# Chromatin Remodeling Drives Immune-Fibroblast Crosstalk in Heart Failure Pathogenesis

**DOI:** 10.1101/2023.01.06.522937

**Authors:** Michael Alexanian, Arun Padmanabhan, Tomohiro Nishino, Joshua G. Travers, Lin Ye, Clara Youngna Lee, Nandhini Sadagopan, Yu Huang, Angelo Pelonero, Kirsten Auclair, Ada Zhu, Barbara Gonzalez Teran, Will Flanigan, Charis Kee-Seon Kim, Koya Lumbao-Conradson, Mauro Costa, Rajan Jain, Israel Charo, Saptarsi M. Haldar, Katherine S. Pollard, Ronald J. Vagnozzi, Timothy A. McKinsey, Pawel F. Przytycki, Deepak Srivastava

## Abstract

Chronic inflammation and tissue fibrosis are common stress responses that worsen organ function, yet the molecular mechanisms governing their crosstalk are poorly understood. In diseased organs, stress-induced changes in gene expression fuel maladaptive cell state transitions and pathological interaction between diverse cellular compartments. Although chronic fibroblast activation worsens dysfunction of lung, liver, kidney, and heart, and exacerbates many cancers, the stress-sensing mechanisms initiating the transcriptional activation of fibroblasts are not well understood. Here, we show that conditional deletion of the transcription co-activator *Brd4* in *Cx3cr1*-positive myeloid cells ameliorates heart failure and is associated with a dramatic reduction in fibroblast activation. Analysis of single-cell chromatin accessibility and BRD4 occupancy *in vivo* in *Cx3cr1*-positive cells identified a large enhancer proximal to *Interleukin-1 beta (Il1b)*, and a series of CRISPR deletions revealed the precise stress-dependent regulatory element that controlled expression of *Il1b* in disease. Secreted IL1B functioned non-cell autonomously to activate a p65/RELA-dependent enhancer near the transcription factor *MEOX1*, resulting in a profibrotic response in human cardiac fibroblasts. *In vivo*, antibody-mediated IL1B neutralization prevented stress-induced expression of *MEOX1*, inhibited fibroblast activation, and improved cardiac function in heart failure. The elucidation of BRD4-dependent crosstalk between a specific immune cell subset and fibroblasts through IL1B provides new therapeutic strategies for heart disease and other disorders of chronic inflammation and maladaptive tissue remodeling.

---

In human disease, stress-induced signaling converges in the nucleus, where the chromatin receives, processes, and amplifies upstream inputs, ultimately leading to altered gene expression, maladaptive cell state transitions and organ dysfunction^1^. In response to stress, chromatin remodeling can alter the expression of secreted factors thus influencing neighboring cells through non-cell autonomous mechanisms and fueling pathological crosstalk between diverse cellular compartments. We previously demonstrated that pharmacological inhibition of bromodomain and extra-terminal domain (BET) proteins, critical mediators of stress-activated chromatin signaling in various diseases^2^, modulates a reversible transcriptional switch that governs fibroblast activation in chronic heart failure through the transcription factor (TF) MEOX1^3^. However, the mechanism by which organ-level stress during heart failure is sensed by cells to trigger downstream fibroblast activation remains unknown. Here, we combined single-cell genomics with dynamic modulation of cell state via small-molecules, cell-type specific genetic perturbation and epigenomic profiling, and CRISPR-based enhancer deletion, to reveal a fundamental and therapeutically relevant mechanism by which pathological cellular crosstalk within a stressed organ drives human disease.

## Correlation of cardiac function and gene expression identifies a stress-activated *Cx3cr1-*expressing monocyte/macrophage subpopulation

To capture the dynamic changes in cell states and identify critical upstream signals that control stress-dependent gene transcription in heart failure pathogenesis, we induced pressure-overload mediated heart failure in mice by performing transverse aortic constriction (TAC) with or without administration of the small molecule BET bromodomain inhibitor JQ1^4^ daily for 30 days (**Fig. 1A**). Consistent with previous observations, BET inhibition exerted a strong protective effect on left ventricular ejection fraction (**Fig. 1B**)^3,5–9^. Given our recent observation that transcriptional changes associated with JQ1 in the cardiomyocyte (CM) compartment are very modest compared to other cardiac cell populations ^3,10^, we performed single cell RNA sequencing (scRNA-seq) in the non-CM compartments of mouse hearts using the 10X Genomics platform (**Fig. 1A**). We captured 41,626 cells from Sham, TAC vehicle-treated (TAC), or TAC JQ1-treated (TAC JQ1) hearts, and identified 20 transcriptional clusters, including a large number of fibroblasts, myeloid and endothelial cells (**Fig. 1C** and **Fig. S1A**).

**Fig. 1.**
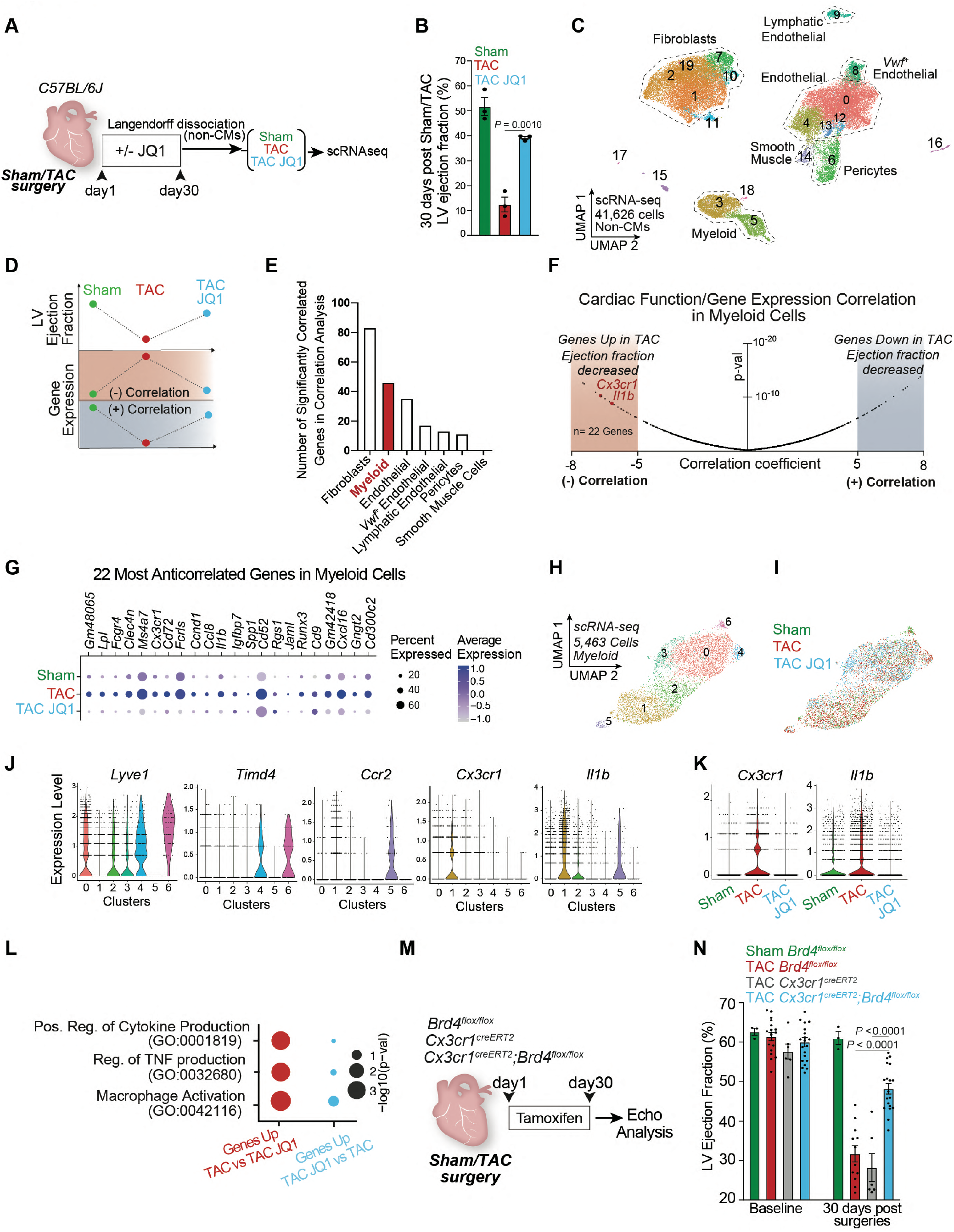
Correlation of cardiac function and gene expression identifies a stress-activated Cx3cr1-expressing macrophage subpopulation that contributes to heart failure pathogenesis. **A.** Schematic of experimental settings of Sham and Transverse Aortic Constriction (TAC) models in C57BL/6J mice with daily dose of the BET inhibitor JQ1 (50mg/kg). **B.** Left Ventricle (LV) ejection fraction quantified by echocardiography. Statistical significance is shown between TAC and TAC JQ1 at day 30 post-TAC. **C.** UMAP plot (scRNA-seq) of non-cardiomyocyte cells colored by cluster. **D.** Schematic of correlation analysis between LV ejection fraction and gene expression highlighting a negative or positive correlation. **E**. Number of strongly positively- and negatively-correlated genes (p-value < 1e^-6^, corresponding to correlation score < −5 and > +5) across the cell population composed by a minimum of 250 cells. **F**. Volcano plot showing correlation coefficients (refer to analysis depicted in Fig. 1D) and corresponding p-values for all the 17,519 genes captured in myeloid cells. Cx3cr1 and Il1b are highlighted in the regions with the most negative correlation coefficients (n=22 genes with score < −5). **G**. Expression Dot Plot of the 22 most-anticorrelated genes in Sham, TAC, and TAC JQ1. **H,I**. UMAP plot (scRNA-seq) of myeloid cells colored by cluster (H) and sample identity (I). **J**. Expression Violin Plots of Lyve1, Timd4, Ccr2, Cx3cr1, and Il1b across myeloid clusters. **K**. Expression Violin Plots of Cx3cr1 and Il1b across samples. **L**. Dot Plot indicating significance (-log10(p-val)) for indicated GO terms in genes upregulated in TAC versus TAC JQ1 (red) or upregulated in TAC JQ1 versus TAC (blue). **M**. Schematic of experimental settings for conditional Brd4 deletion in Cx3cr1^Pos^ cells. **N**. LV ejection fraction quantified by echocardiography in Cx3cr1^CreERT2^ (CRE control) and Brd4^flox/flox^ and Cx3cr1^CreERT2^;Brd4^flox/flox^ littermates. **B**,**N** Data are mean ± s.e.m. One-way ANOVA followed by Tukey post hoc test.

To identify dynamically regulated cell states during stress and BET inhibition we developed a model in which we correlated the changes in systolic heart function associated with TAC and JQ1 with changes in gene expression. Genes were defined as having a negative correlation if their expression was anticorrelated with heart function (i.e., upregulated when cardiac function decreased and downregulated when cardiac function increased), or a positive correlation in the inverse scenario (**Fig. 1D**). We performed this analysis in each cellular compartment that was composed of at least 250 cells, namely fibroblasts, myeloid cells, endothelial cells, von Willebrand factor (*VWf*)-positive endothelial cells, lymphatic endothelial cells, pericytes, and smooth muscle cells (SMCs). As expected^3^, fibroblasts had the highest enrichment of genes whose expression significantly correlated (either negatively or positively) with systolic cardiac function (**Fig. 1E**). Interestingly, myeloid cells had the next highest number of correlated genes, indicating a high degree of transcriptional plasticity in this compartment in response to stress and BET inhibition (**Fig. 1E**). Given the ability of myeloid cells to act as noncell autonomous amplifiers of the stress-response through secreted factors^11–13^, we further investigated changes in this cell type.

As BET proteins are transcriptional co-activators and their inhibition is associated with protection in heart failure models^5–7^, we focused on genes in myeloid cells that were upregulated in disease and downregulated with BET inhibition. We ranked all genes expressed in the myeloid compartment based on their correlation score using a stringent statistical threshold (*p*-value < 1e^-6^, corresponding to a correlation score < −5) and identified a total of 22 genes whose expression was strongly anti-correlated with cardiac function (**Fig. 1F**, left, highlighted in red). Among these genes were cytokines (*Il1b*), chemokines (*Ccl8, Ccl12*), chemokine receptors (*Cx3cr1*), cluster of differentiation (CD) proteins (*Cd9, Cd52*, and *Cd72*) and the TF *Runx3* (**Fig. 1G**). Sub-clustering of the myeloid population identified 7 clusters distributed over 5,463 cells (**Fig. 1H,I**). Expression of the resident macrophage marker^14,15^ *Lyve1* was enriched in clusters 0, 2, 3, 4, and 6, with *Timd4* marking only clusters 4 and 6 (**Fig. 1J** and **Fig. S1B-D**). Clusters 1 and 5 had high expression of *Ccr2*, a marker of monocyte-derived macrophages that infiltrate from the circulation^16–18^ (**Fig. 1J** and **Fig. S1B-D**). *Cx3cr1*, which marks both monocyte-derived and resident macrophages^19–21^, was particularly enriched in cluster 1, together with proinflammatory genes such as *Il1b* (**Fig. 1J**). Consistent with *Cx3cr1* and *Il1b* expression being negatively correlated with systolic heart function (**Fig. 1F**), both genes were increased in TAC compared to Sham, and downregulated by JQ1 treatment (**Fig. 1K**). *Cx3cr1* and *Il1b* were co-expressed in cluster 1 (**Fig. 1J**), and notably this cluster was enriched in cells from TAC, and depleted in Sham and TAC JQ1 conditions (**Fig. S1E**). Differential expression (DE) analysis revealed that the most enriched gene ontology (GO) terms of the genes that were significantly upregulated in TAC versus TAC JQ1 were associated to proinflammatory mechanisms such as regulation of cytokine and TNF production and macrophage activation (**Fig. 1L**).

## BRD4 in *Cx3cr1*-expressing cells contributes to heart failure pathogenesis

To test the functional significance of BET-dependent chromatin signaling in *Cx3cr1-expressing* monocytes and macrophages, we genetically deleted the BET family member BRD4 in *Cx3cr1*-expressing cells by crossing *Brd4^flox/flox^* mice^10^ with a mouse in which tamoxifen-inducible Cre recombinase is directed by the *Cx3cr1* promoter region^19^ (*Cx3cr1-CreERT2*) (**Fig. 1M**). While this approach leaves other BET family members intact, we previously showed the specific importance of BRD4 in aspects of heart failure pathogenesis^3,22^. Mice were subjected to TAC and 5 days of continuous tamoxifen treatment via intraperitoneal injection followed by an injection every other day until day 30 post-TAC, a regimen that would delete *Brd4* in both resident and monocyte-derived *Cx3cr1*-expressing macrophages^14^. To confirm cell-type specific loss of *Brd4* we performed Langendorff perfusion, isolated CMs and then flow sorted *Cx3cr1*-expressing cells (CX3CR1^Pos^, CD45^Pos^), fibroblasts (CD45^Neg^, CD31^Neg^, mEF-SK4^Pos^), and endothelial cells (CD45^Neg^, CD31^Pos^, mEF-SK4^Neg^). *Brd4* mRNA levels were significantly reduced in *Cx3cr1^CreERT2^;Brd4^flox/flox^* versus *Brd4^flox/flox^* animals only in *Cx3cr1^pos^* cells, while *Brd4* expression was not affected in CMs, fibroblasts or endothelial cells (**Fig. S1F**). Cardiac function was evaluated by transthoracic echocardiography at baseline and 30 days after TAC. When compared with littermate controls (*Cx3cr1^CreERT2^* or *Brd4^flox/flox^*) undergoing the same experimental manipulations, *Cx3cr1^CreERT2^;Brd4^flox/flox^* mice demonstrated improved cardiac contractile function (**Fig. 1N**). This suggests *Brd4* function in *Cx3cr1*-positive cells contributes substantially to stress-dependent cardiac dysfunction.

## BRD4 triggers a proinflammatory transcriptional response in heart failure pathogenesis

To determine the transcriptional consequences of *Brd4* deletion in the immune cell compartment, we dissociated cardiac ventricle tissue into single cell preparations and performed scRNA-seq on sorted CD45-positive cells from *Brd4^flox/flox^* mice after TAC or Sham surgery, and from Cx3cr1^CreERT2^;Brd4^flox/flox^ mice after TAC (TAC-*Brd4*KO) **(Fig. 2A**). We processed 10,450 immune cells and identified 16 transcriptomic clusters. Most of the cardiac CD45-positive cells were grouped in a large and heterogeneous population of monocytes/macrophages (clusters 0 to 10 excluding 7, **Fig. S 2A-C**), while B cells (cluster 14), T cells (cluster 11) and neutrophils (cluster 12) were fewer in number (**Fig. S2A,C**). Upon re-clustering of only the monocyte/macrophage population (**Fig. 2B**), it was evident that expression of the resident macrophage markers *Lyve1* and *Timd4* was reduced in cluster 4, which together with high expression of *Ccr2* and *Cx3cr1* suggested that this cluster was likely composed of monocytes that had infiltrated the heart following TAC (**Fig. 2C,D**). Clusters 3 and 4 were mostly composed of cells from Sham and TAC samples and not TAC-*Brd4*KO (**Fig. 2E**). We calculated the percentage of cells from Sham, TAC and TAC-*Brd4*KO samples belonging to each of the 11 clusters and found that cluster 4 was associated with an expansion from Sham to TAC that was strongly reduced upon *Brd4* deletion (**Fig. S2D**). GO analysis of the genes driving cluster 4 identified as top enriched terms processes related to the pro-inflammatory response, such as cellular response to lipopolysaccharide (LPS) and positive regulation of cytokine production (**Fig. 2F**). Among the top ten cluster 4 marker genes were several encoding the major histocompatibility complex (MHC) class 2 glycoproteins (also present in cluster 3) and genes such as *Il1b* and *Ccr2*, both of which were very specific to cluster 4 (**Fig. 2G**). We performed DE analysis in the monocyte/macrophage population between samples (**Fig. S2E**), and applied a strict statistical threshold to focus on genes that were strongly triggered by stress (upregulated TAC versus Sham, LogFC>0.5; FDR<0.05, total of 48) and decreased by *Brd4* deletion in *Cx3cr1*-positive cells (downregulated in TAC-*Brd4*KO versus TAC, LogFC>0.5; FDR<0.05, total of 66). This analysis identified 13 genes including *Rgs1, Il1b, Cd52, Cd72*, and *Cx3cr1* (**Fig. 2H**). The expression of the proinflammatory cytokine *Il1b* was particularly interesting as it was enriched in cluster 4 (**Fig. 2I,J**) and very sensitive to both cardiac stress and *Brd4* deletion in *Cx3cr1*-positive cells (**Fig. 2K**).

**Fig. 2.**
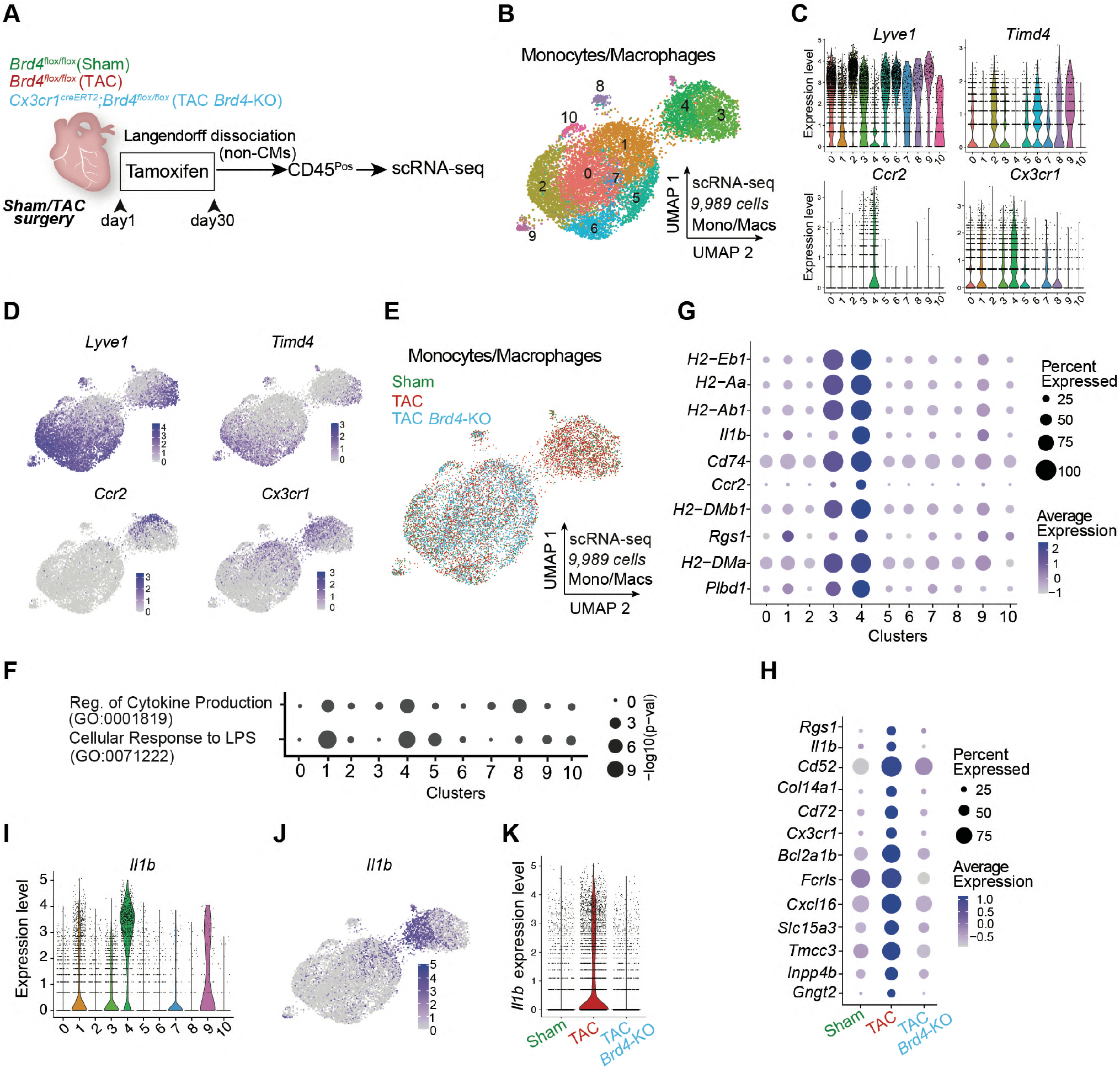
BRD4 in Cx3cr1-expressing macrophages triggers a proinflammatory transcriptional response in heart failure pathogenesis. **A.** Schematic of experimental settings for scRNA-seq in sorted CD45^Pos^ cells from hearts of Brd4^flox/flox^ (TAC) and Cx3cr1^CreERT2^;Brd4^flox/flox^ (TAC Brd4-KO) littermates. **B.** UMAP plot (scRNA-seq) of monocytes/macrophages colored by cluster. **C.** Expression Violin Plots of Lyve1, Timd4, Ccr2, and Cx3cr1 across monocyte/macrophage clusters. **D.** Expression Feature Plots (scRNA-seq) of Lyve1, Timd4, Ccr2, and Cx3cr1 across monocytes and macrophages. **E**. UMAP plot (scRNA-seq) of monocytes/macrophages colored by sample identity. **F**. Dot Plot indicating significance (-log10(p-val)) for indicated GO terms across all monocyte/macrophage clusters. **G**. Expression Dot Plot of the 10 top gene markers in cluster 4 across all monocyte/macrophage clusters. **H**. Expression Dot Plots (scRNA-seq) of the 13 genes upregulated TAC versus Sham and downregulated in TAC-Brd4KO versus TAC in monocytes/macrophages across samples (LogFC>0.5 and FDR<0.5). **I**. Expression Violin Plots of Il1b across monocyte/macrophage clusters. **J**. Expression Feature Plot (scRNA-seq) of Il1b in monocytes and macrophages. **K**. Expression Violin Plot of Il1b in monocyte/macrophage across samples.

## BRD4 in *Cx3cr7*-expressing cells function non-cell autonomously to activate cardiac fibroblasts in stress

Since many cytokines and chemokines were dysregulated in immune cells upon *Brd4* deletion in *Cx3cr1*-positive cells, we investigated the non-cell autonomous consequences in other cell types, including CMs, in the setting of cardiac stress. We harvested hearts from TAC or TAC-*Brd*4KO animals, isolated nuclei, and performed single nuclei RNA-seq (snRNA-seq) to capture the transcriptional landscape of all cardiac cells (**Fig. 3A**). A group of 28,107 nuclei encompassing all the major cardiac cell populations was processed into 17 transcriptomic clusters (**Fig. 3B**, **Fig. S3A**). The cardiomyocyte population was less represented than expected, likely due to the process of nuclei isolation. The UMAP suggested striking differences in myeloid cells upon *Brd4* deletion in *Cx3cr1*-positive cells as expected, but also indicated similar changes in the cardiac fibroblast population (**Fig. 3C**). DE analysis between the TAC and TAC-*Brd4*KO conditions in each of the major cardiac cell populations (LogFC>0.125; FDR<0.05) also demonstrated that a similar number of genes were dysregulated in myeloid cells (n=351) and fibroblasts (n=310), with far fewer in other cell types (**Fig. 3D**). Since *Cx3cr1* is not expressed in fibroblasts^23^, and thus *Brd4* is not deleted in this cell population, the gene expression changes in fibroblasts are secondary to BRD4 loss in *Cx3Cr1*-positive cells and therefore reflect a non-cell autonomous effect.

**Fig. 3.**
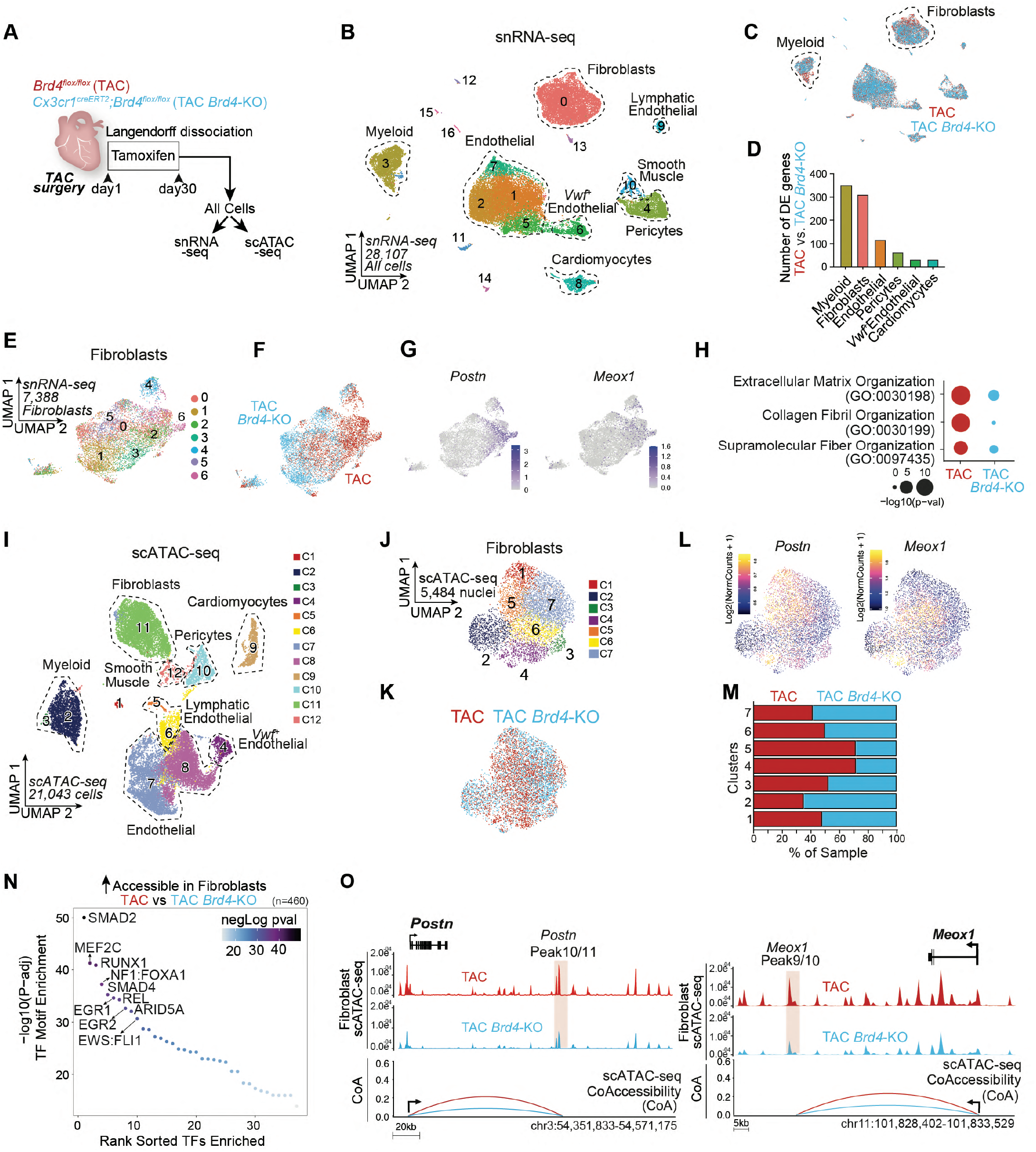
BRD4 in Cx3cr1-expressing cells functions non-cell autonomously to activate cardiac fibroblasts under stress. **A.** Schematic of experimental settings for single nuclei RNA-seq and ATAC-seq from hearts of Brd4^flox/flox^ (TAC) and Cx3cr1^CreERT2^;Brd4^flox/flox^ (TAC Brd4-KO) littermates. **B,C.** UMAP plot (snRNA-seq) of nuclei from cardiac tissue colored by cluster (B) and sample identity (C). **D.** Number of DE genes between TAC and TAC Brd4-KO across cell populations with at least 800 cells (LogFC>0.125; FDR<0.05). **E,F**. UMAP plot (snRNA-seq) of fibroblasts colored by cluster (E) and sample identity (F). **G**. Expression Feature Plot (snRNA-seq) of Postn and Meox1 in fibroblasts. **H**. Dot Plot indicating significance (-log10(p-val)) for indicated GO terms in genes upregulated in TAC versus TAC JQ1 (red) or upregulated in TAC JQ1 versus TAC (blue) in fibroblasts. **I**. UMAP plot (scATAC-seq) of all cardiac cells colored by cluster. **J, K**. UMAP plot (scATAC-seq) of fibroblasts colored by cluster (J) and sample identity (K). **L**. Chromatin accessibility gene score of Postn and Meox1 in fibroblasts. **M**. Sample distribution within clusters in fibroblasts. **N**. TF motif enrichment of fibroblast chromatin regions more accessible in TAC versus TAC Brd4-KO. Top 10 motifs are highlighted. **O**. Fibroblast chromatin accessibility in TAC and TAC Brd4-KO at the Postn and Meox1 locus. The signal at essential regulatory elements is highlighted in red together with Co-Accessibility with the respective gene promoter.

Within the fibroblast population, we identified 7 separate transcriptional clusters encompassing 7,388 cells (**Fig. 3E** and **Fig. S3B**), among which TAC and TAC-*Brd4*KO fibroblasts clustered separately (**Fig. 3E,F**). Cluster 6 was associated with specific expression of genes involved in fibroblast activation such as *Postn, Meox1, Comp, Cilp, Mfap4*, and *Thbs4* (**Fig. 3G**, **Fig. S3B-D**), and was characterized by a depletion of TAC-*Brd4*KO fibroblasts (**Fig. S3E**). GO analysis of the genes that were significantly upregulated in TAC versus TAC-*Brd4*KO (n=198) (**Fig. S3F**, the top 50 are shown) was enriched for terms related to profibrotic gene programs such as extracellular matrix and collagen fibril organization (**Fig. 3H**). In contrast, these terms were less represented in the genes upregulated in TAC-*Brd4*KO versus TAC (n=112) (**Fig. 3H**), which were instead enriched with terms associated with the regulation of vasculature development, angiogenesis, and response to VEGF (**Fig. S3G**).

To understand the changes in chromatin states in cardiac tissue triggered by the deletion of *Brd4* in *Cx3cr1*-expressing cells, we performed single cell Assay for Transposase-Accessible Chromatin sequencing (scATAC-seq) from the same hearts used for snRNA-seq (**Fig. 3A**). As expected, scATAC-seq annotated all the major cell populations identified in the snRNA-seq analysis (**Fig. 3I** and **Fig. S4A**).

Given the profound changes in gene expression identified in fibroblasts (**Fig. 3D**), we focused on the corresponding changes in chromatin accessibility in this population. Subsetting the fibroblast cells identified 7 clusters (**Fig. 3J,K**), and gene scores (a measure of chromatin accessibility within and around the gene body) suggested that clusters 4 and 5 were particularly enriched with signals from *Postn* and *Meox1*, as well as other markers of the activated fibroblast state (**Fig. 3L** and **Fig. S4B,C**). Assessment of sample distributions within the scATAC-seq clusters identified clusters 4 and 5 as the most depleted in TAC-*Brd4*KO samples (**Fig. 3M**). We then ran differential accessibility analysis between TAC and TAC-*Brd4*KO to identify sites that lost chromatin accessibility in fibroblasts when *Brd4* was deleted in *Cx3cr1*-expressing cells (n=460). Motif analysis of these regions revealed enrichment for sites predicted to bind TFs known to be critical regulators of the profibrotic response in heart failure, such as SMADs^24^, as well as motifs for RUNX1 and REL (**Fig. 3N**). While the function of the latter TFs in governing the proinflammatory response is established^24^, their role in controlling the profibrotic response in heart failure is not well understood. We also examined accessibility of critical stress-induced enhancers of *Postn* and *Meox1* that we described previously^3^. We found that the large distal elements encompassing the *Postn* and *Meox1* locus displayed a marked reduction of chromatin accessibility in TAC-*Brd4*KO fibroblasts, including the specific regulatory elements *Postn Peak 11* and *Meox1 Peak 9/10* that we have previously shown to be essential for the stress-induced activation of their cognate genes (**Fig. 3O**, regions highlighted in red)^3^.

## Chromatin accessibility and BRD4 occupancy *in vivo* identify the precise regulatory elements that control stress-dependent activation of *Il1b*

Having found that a BRD4-dependent transcriptional event in *Cx3cr1*-expressing cells functions non-cell autonomously to activate cardiac fibroblasts under stress, we performed scATAC-seq in the sorted CD45-positive cells from Sham, TAC, and TAC-*Brd4*KO animals to determine how *Brd4* deletion in *Cx3cr1*-positive cells directly affects chromatin accessibility in the immune compartment (**Fig. 4A**). We identified 21 clusters, and gene score signals of *C1qa, Cd163*, and *Cd14* identified the clusters encompassing the monocyte/macrophage population, which was also accessible in the *Cx3cr1* locus (**Fig. S5A,B**). Subsetting the monocyte/macrophage population identified 8 distinct clusters (**Fig. 4B,C**). Chromatin accessibility of *Timd4* and *Lyve1* were high in cluster 3, consistent with resident macrophages, while cluster 4 displayed strong signal for the monocyte-derived macrophage marker *Ccr2* (**Fig. 4D**). Interestingly, *Cx3cr1* accessibility was almost specific to cluster 4, as was accessibility of *Il1b*, one of the most dynamically regulated genes in the scRNA-seq analysis (**Fig. 4D**). The *Cx3cr1/Il1b* accessible cluster 4 was more represented by cells from the TAC condition compared to Sham, and *Brd4* deletion was associated with a decrease in its composition (**Fig. S5C,D**). GO analysis of the accessible chromatin regions driving cluster 4 identified proinflammatory processes such as regulation of cytokine production as top enriched gene programs (**Fig. 4E**). TF motif analysis of these same regions, compared to the *Timd4/Lyve1* accessible cluster 3, found REL, IKZF1, and RUNX2 at the top 3 most enriched motifs (**Fig. 4F**).

**Fig. 4.**
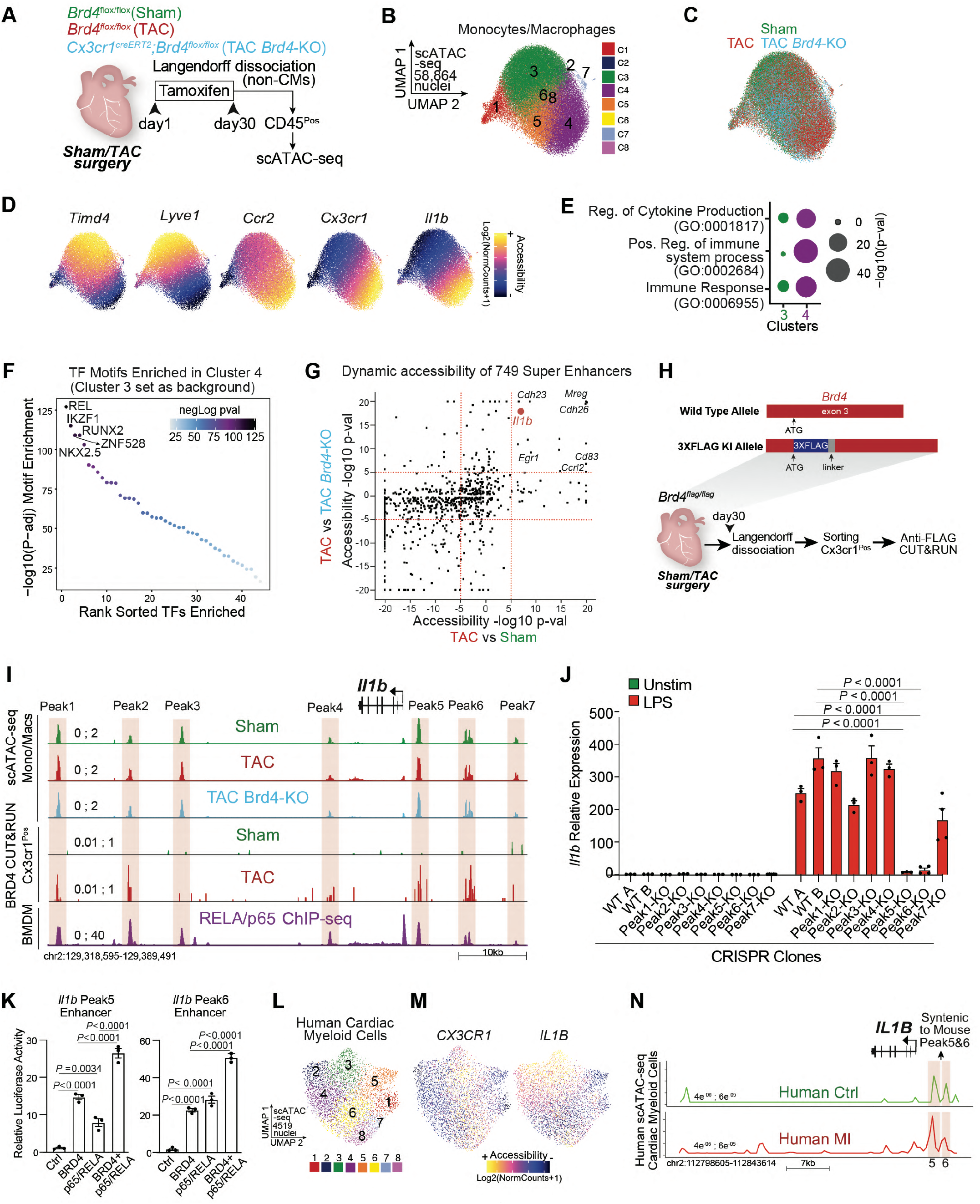
Chromatin accessibility and BRD4 occupancy in vivo identify the precise regulatory elements that control stress-dependent activation of Il1b. **A.** Schematic of experimental settings for scATAC-seq in sorted CD45^Pos^ cells from hearts of Brd4^flox/flox^ (TAC) and Cx3cr1^CreERT2^;Brd4^flox/flox^ (TAC Brd4-KO) littermates. **B,C.** UMAP plot (scATAC-seq) of monocytes/macrophages colored by cluster (B) and sample identity (C). **D.** Chromatin accessibility gene score of Timd4, Lyve1, Ccr2, Cx3cr1, and Il1b in monocytes/macrophages. **E**. Dot Plot indicating significance (-log10(p-val)) for indicated GO terms in genes driving monocyte/macrophage clusters 3 and 4. **F**. TF motif enrichment of monocyte/macrophage chromatin regions in cluster 4 (cluster 3 was used as background). Top 5 motifs are highlighted. **G**. Chromatin Accessibility changes between TAC and Sham (x axis) and TAC and TAC Brd4-KO (y axis) for the 749 super enhancers identified in monocytes/macrophages. Dotted red lines at +5 and −5 in both axes are indicated. Genes proximal to candidate enhancers are highlighted in top right quadrant. **H**. Targeting strategy to generate Brd4^flag/flag^ animals (top) and schematic of experimental settings for anti-FLAG CUT&RUN performed on Cx3cr1^Pos^ cells from Brd4^flag/flag^ Sham and TAC mice (bottom). **I**. Il1b locus showing from top to bottom: scATAC-seq mean normalized accessibility across monocytes/macrophages in Sham, TAC and TAC Brd4-KO; anti-FLAG CUT&RUN signal in Cx3cr1^Pos^ cells from Brd4^flag/flag^ Sham and TAC mice; RELA/p65 ChIP-seq on BMDMs. 7 regions (Peak1 to 7) of high chromatin accessibility are highlighted in red. **J**. Il1b expression by qPCR in Unstimulated or LPS (1ng/ml) treated RA W 264-7 macrophages in CRISPR WT and Peak1 to Peak7 KO clones. Significance is shown between the 2 WT clones and Il-1b Peak5-KO and Il-1b Peak6-KO clones. **K**. Transcriptional activity by luciferase assay of Il-1b Peak5 and Peak6 enhancer regions in the presence of BRD4 and p65/RELA. **L**.UMAP plot (scATAC-seq) of human cardiac myeloid cells^28^ colored by cluster. **M**. Chromatin accessibility gene score of CX3CR1 and IL1B in human cardiac myeloid cells. **N**. Human IL1B locus showing scATAC-seq mean normalized accessibility across human cardiac myeloid cells in controls (green) and patients with myocardial infarction (red). Red boxes with the same height of the y axis are placed on peak 5 and 6. **J**,**K**, Data are mean ± s.e.m. Two-way (J) and One-way (K) ANOVA followed by Tukey post hoc test.

To determine which distal elements in the monocyte/macrophage population were most sensitive to stress and *Cx3cr1*-specific *Brd4* deletion, we assembled a catalog of 749 large enhancers (also known as super-enhancers)^25^ using scATAC-seq data from diseased hearts (TAC) (**Fig. S6A**). We then constructed a model that identified distal regions with highly dynamic changes across conditions, and scored the enhancers by their degree of changes in TAC versus Sham, and in TAC versus TAC-*Brd4*KO (**Fig. 4G**). We were particularly interested in regions that opened with cardiac stress and had decreased accessibility upon *Brd4* deletion (**Fig. 4G**, top right quadrant). Among the 15 most affected regions were DNA elements proximal to genes such as *Il1b, Mreg, Cdh26, Cdh23, Cd83, Ccrl2*, and *Egr1*. To determine which of these loci were directly bound by BRD4 *in vivo* specifically in *Cx3cr1*-expressing cells, we generated a mouse in which a 3x-FLAG epitope tag was knocked-in into the N-terminus of BRD4 (**Fig. S6B-D**) and performed the highly sensitive anti-FLAG CUT&RUN (Cleavage Under Targets & Release Using Nuclease) on *Cx3cr1*-positive sorted cells from Sham or TAC mice (**Fig. 4H** and **Fig. S6E**). Among the 15 dynamic regions that exhibited exquisite dynamic chromatin accessibility across conditions (**Fig. 4G**, top right quadrant), 10 were associated with higher binding of BRD4 in TAC including the large enhancers proximal to *Il1b, Mreg, Cdh26*, and *Cd83* (**Fig. S6F**).

Notably, one of the most dynamic BRD4-bound distal elements in stress and upon genetic deletion of *Brd4* was proximal to *Il1b* (**Fig. 4G**). *Il1b* was amongst the most negatively correlated genes in the analysis between cardiac function and gene expression in myeloid cells following stress and BET inhibition (**Fig. 1F**). Moreover, *Il1b* was one of the most sensitive genes to *Brd4* deletion in *Cx3cr1*-expressing cells, suggesting direct regulation by BRD4 (**Fig. 2H,K**). Importantly, *Il1b* encodes a secreted factor (IL1B) and thus has the ability to influence other cell states and modulate pathological cellular crosstalk in heart disease via paracrine signaling. To identify the functionally relevant stress-dependent distal elements within the large 71kb region in the *Il1b* locus, we examined the 3 accessible peak regions upstream and the 4 peaks downstream of *Il1b* (**Fig. 4I**) as determined by scATAC-seq. Overlaying BRD4 CUT&RUN results from the sorted *Cx3cr1*-positive cells with the scATAC-seq data demonstrated that upon cardiac stress, peaks 1, 2, 5, and 6 were characterized by strong recruitment of BRD4 (**Fig. 4I**).

Given that p65/RELA regulates *Il1b* expression in other cell types^26^, and that REL was the most enriched TF motif in the accessible DNA regions of *Cx3cr1*-positive cells (**Fig. 4F**), we used publicly available data in bone marrow-derived macrophages (BMDMs)^27^ to find that p65/RELA directly bound all the peaks identified within the *Il1b* enhancer (**Fig. 4I**). To systematically identify which regions, if any, were functional *cis*-regulatory elements, we used CRISPR-Cas9 in RAW 264-7 macrophages to target each individual enhancer peak within the *Il1b* locus and generated 7 different clones, each harboring a specific deletion for one of the 7 peaks (**Fig. S 6G-I)**. Lipopolysaccharides (LPS) triggered a strong upregulation of *Il1b* gene expression in the control WT clones (exposed to CRISPR machinery and non-targeting gRNAs) (**Fig. 4J**). Notably, while most of the CRISPR deletions did not affect LPS-induced *Il1b* induction significantly, deletion of peaks 5 and 6 completely abolished stress-induced expression (**Fig. 4J**). Given the colocalization of BRD4 and p65/RELA at the *Il1b* Peak5 and 6 enhancers, we hypothesized that these factors may functionally coregulate these regulatory elements. To test this, we separately cloned the *Il1b* Peak5 and 6 enhancers into a luciferase reporter, and demonstrated additive coregulation by BRD4 and p65/RELA in controlling the transcriptional activity of these regions (**Fig. 4K**).

Finally, we tested whether we could identify corresponding changes in chromatin states in human cardiac myeloid cells. Single-cell ATAC-seq data from the human adult heart^28^ identified 8 distinct clusters in myeloid cells (**Fig. 4L**), with cluster 3 being associated with strong accessibility for both *CX3CR1* and *IL1B* (**Fig. 4M**), while clusters 5,6 and 8 were associated with high accessibility for *TIMD4* and *LYVE1* (**Fig. S5E**). GO analysis of the accessible chromatin regions driving human cluster 3 highlighted enrichment of proinflammatory processes as compared to other clusters (**Fig. S5F**). We then focused on the human *IL1B* locus and found that the syntenic regions of mouse *Il1b* Peak5 and 6 enhancers were the most accessible regions within the locus, suggesting these regions might be critical for regulating *IL1B* expression in humans (**Fig. 4N**). Importantly, human *IL1B* Peak5 was associated with increased accessibility in individuals with myocardial infarction as compared to controls, highlighting dynamic regulation of this specific region in disease (**Fig. 4N**).

## IL1B promotes profibrotic function and activates *MEOX1* through a p65/RELA-dependent enhancer

Given the highly dynamic regulation of *Il1b* in the *Cx3cr1*-positive population, and the corresponding changes in transcription and chromatin accessibility in the cardiac fibroblasts upon *Brd4* deletion in *Cx3cr1*-expressing cells, we hypothesized that IL1B could be a critical signal secreted from the *Cx3cr1*-positive cells under the direct control of BRD4 to activate the pathological fibroblast stress response in heart failure. Importantly, fibroblasts express *Il1r1*, the receptor to which IL1B binds to propagate its signal (**Fig. S7A,B**)^29^. We used human induced pluripotent stem cells (iPS) differentiated into cardiac fibroblasts^30^ (**Fig. S7C**) to test whether IL1B could modulate the profibrotic response, particularly in the context of its effect in combination with TGFB, a master signaling protein for fibroblast activation in a wide variety of human disorders including heart disease^31–33^. IL1B led to a significant increase in the formation of α-smooth muscle (αSMA)-positive stress fibers, and importantly demonstrated an increased signal when used in combination with TGFB compared to TGFB alone (**Fig. S7D**). We then investigated whether IL1B could affect cardiac fibroblast contractility, a functional hallmark of fibroblast activation in disease pathogenesis. Indeed, IL1B in combination with TGFB triggered increased collagen-contraction compared to TGFB alone (**Fig. S7E**). Interestingly, treatment of IL1B alone over time was able to reach the level of contractility achieved by TGFB alone (**Fig. S7E**). To more directly test if secreted IL1B can function in a paracrine fashion to activate cardiac fibroblasts, we treated BMDMs with LPS, which notably activates a strong proinflammatory response including IL1B secretion (**Fig. S7F**). We collected the medium and treated it for 2 hours with 10 ug/ml of either an isotype control IgG antibody, or an anti-IL1B neutralizing antibody, and then added the conditioned medium onto iPS-derived cardiac fibroblasts plated on collagen pads (**Fig. 5A**). While the medium treated with LPS and IgG antibody was able to strongly increase fibroblast contractility compared to medium from unstimulated BMDMs, incubation with the neutralizing IL1B significantly attenuated collagengel contraction, demonstrating that IL1B endogenously produced by myeloid cells can function in a paracrine fashion to modulate the contractile phenotype of activated fibroblasts (**Fig. 5A**).

**Fig. 5.**
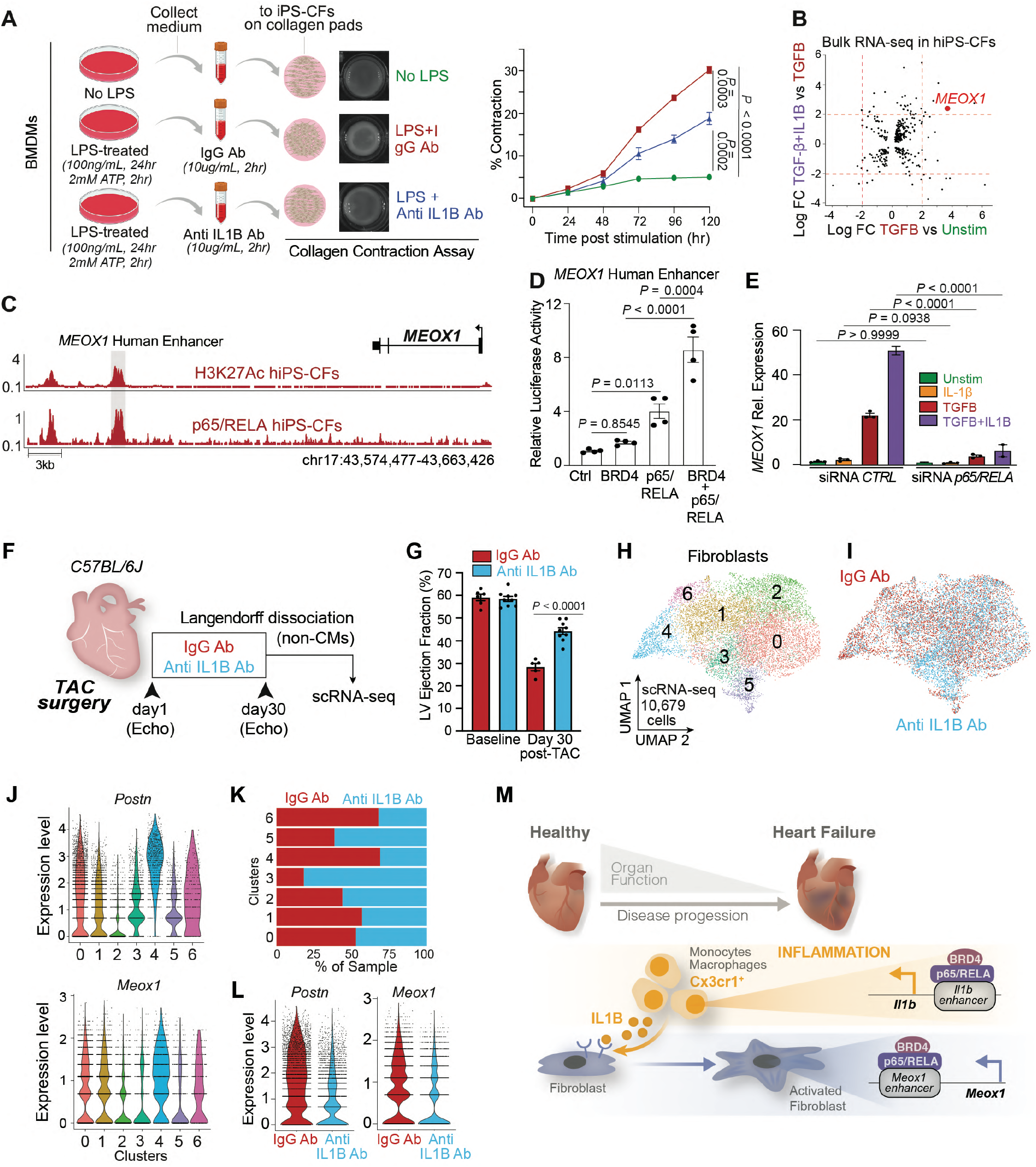
IL1B neutralization improves cardiac function in heart failure and prevents fibroblast activation. **A.** Schematic representation of experiment (left): medium from unstimulated or LPS-treated BMDM is treated with either IgG isotype control or anti-IL1B antibodies, then the conditioned medium is added onto the iPS-CFs and collagen contractility over 120 hours is measured (right). **B**. Bulk RNA-seq in iPS-CFs. LogFC between indicated conditions is shown. Dotted red lines at +2 and −2 in both axes are indicated. MEOX1 gene in the top-right quadrant is highlighted. **C.** Human MEOX1 locus showing H3K27Ac and p65/RELA in unstimulated iPS-CFs. The syntenic region of the mouse Meox1 Peak9/10 regulatory element is highlighted. **D.** Transcriptional activity by luciferase assay of MEOX1 human enhancer region (highlighted in Fig. 5C) in the presence of BRD4 and p65/RELA. **E**. MEOX1 expression by qPCR in Unstimulated iPS-CFs or treated with IL1B, TGF-B, or TGFB+IL1B with control or p65/RELA-targeting siRNAs. **F**. Schematic of experimental settings in TAC C57BL/6J mice treated with either 500ug of IgG isotype control and anti-IL1B antibodies injected intraperitoneally every 3 days. Non-CM cells were processed at day 30 for scRNA-seq. **G**. LV ejection fraction quantified by echocardiography. Statistical significance is shown between IgG and Anti-IL1B group 30 days post-TAC. **H**,**I**. UMAP plot (scRNA-seq) of fibroblasts colored by cluster (H) and sample identity (I). **J**. Expression Violin Plots of Postn and Meox1 across fibroblasts clusters. **K**. Sample distribution within clusters in fibroblasts. **L**. Expression Violin Plot of Postn and Meox1 in fibroblasts across samples. M. Working model depicting the molecular mechanisms regulating the crosstalk between Cx3cr1-positive cells and activated fibroblasts through IL1B and MEOX1. **A**,**D,E** Data are mean ± s.e.m. Two-way (A) and One-way (D,E) ANOVA followed by Tukey post hoc test.

We then investigated the molecular mechanisms involved in the profibrotic response triggered by IL1B, particularly in the context of its effect in combination with TGFB. We performed bulk RNA-seq and DE analysis using a strict statistical threshold (LogFC > 2) to identify genes that were very sensitive to TGFB stimulation and that had increased expression when TGFB and IL1B were used in combination (**Fig. 5B**). This analysis identified a small group of genes such as *SERPINA9, EGR2, LRRC15^34^* and *MEOX1*, a TF that we have recently demonstrated to be a critical regulator of the cell state transition between quiescent and activated cardiac fibroblasts associated with cardiac dysfunction^3^. Given our observation that REL was one of the most enriched TF motifs in fibroblast chromatin regions that were opened in TAC and closed when *Brd4* was deleted in *Cx3cr1*-expressing cells (where *Il1b* is downregulated) (**Fig. 3N**), and that the TF p65/RELA is notably activated by IL1B signaling^35^, we investigated whether p65/RELA was directly involved in *MEOX1* regulation. We performed chromatin immunoprecipitation followed by sequencing (ChIP-seq) for H3K27Ac, which marks active chromatin regions, in human iPS-cardiac fibroblasts. Within the human *MEOX1* locus, we found that the region associated with strong enrichment of H3K27Ac was the syntenic element of the mouse *Meox1 Peak9/10* enhancer^3^ (**Fig. 5C**). We found several motifs for p65/RELA within this 2.7kb distal region (**Fig. S7G**), and p65/RELA ChIP-seq on human cardiac fibroblasts revealed direct chromatin binding at the *MEOX1* human enhancer (**Fig. 5C**). As our previous work demonstrated that *Meox1* transcription is sensitive to BET inhibition^3^, we tested whether BRD4 and p65/RELA functionally coregulate the human *MEOX1* regulatory element. We cloned this *cis*-element into a luciferase reporter and found synergistic coregulation by BRD4 and p65/RELA on the *MEOX1* enhancer (**Fig. 5D**). To test whether *MEOX1* was sensitive to *p65/RELA* depletion, we used an siRNA to downregulate *p65/RELA* (**Fig. S7H**), and found that its knockdown abolished TGFB/IL1B-induced *MEOX1* activation (**Fig. 5E**).

## Antibody-mediated IL-1B neutralization prevents stress-induced fibroblast activation and improves cardiac function in heart failure

Finally, we tested whether blocking IL1B *in vivo* could decrease fibroblast activation and the related cardiac dysfunction in response to pressure-overload stress. We performed TAC surgeries in C57BL6/J mice, and treated them every 3 days with either IgG control or IL1B neutralizing antibodies for 30 days (**Fig. 5F**). Remarkably, antibody-mediated IL1B neutralization exerted a protective effect and significantly increased systolic heart function compared to the isotype control (**Fig. 5G**), achieving a similar level of protection as seen with *Brd4* deletion in *Cx3cr1*-positive cells (**Fig. 1N**). We harvested the hearts from IgG and anti-IL1B treated animals, performed Langendorff perfusion followed by scRNA-seq in the non-cardiomyocyte cell compartment (**Fig. 5F**). We captured 24,279 cells in 14 transcriptomic clusters, encompassing a large population of myeloid, endothelial, and fibroblast cells (**Fig. S8A,B**). Given our interest in the fibroblast population, we subsetted this population for further analysis (**Fig. 5H,I**) and specifically examined expression of genes associated with fibroblast activation and profibrotic function. Among the 7 different clusters identified, clusters 4 and 6 were enriched in fibroblast stress genes such as *Postn, Meox1, Comp, Cilp, Mfap4*, and *Thbs4* (the latter of which was specifically expressed in cluster 4) (**Fig. 5J** and **Fig. S8C,D**). We then calculated sample distribution and found that clusters 4 and 6 were depleted of fibroblasts from anti-IL1B treated animals (**Fig. 5K**). In contrast, cluster 3 was predominantly from anti-IL1B treated hearts and was not enriched for *Postn* or *Meox1*. Indeed, the expression of *Postn* and *Meox1*, and of many other activated fibroblast markers, was strongly downregulated with antibody-mediated IL1B neutralization (**Fig. 5L** and **Fig. S8E,F**). GO analysis of the genes that were significantly upregulated in fibroblasts in IgG versus anti-IL1B treated animals (n=469) identified as top enriched biological processes profibrotic gene programs such as extracellular matrix and collagen fibril organization (**Fig. S8G**). Similar to changes in the fibroblast transcriptome when *Brd4* was deleted in *Cx3cr1*-expressing cells (**Fig. S3G**), the GO terms of the genes upregulated with IL1B neutralization (n=167) were enriched with gene-programs associated with angiogenesis and response to VEGF (**Fig. S8H**), suggesting common fibroblast transcriptional changes associated with cell-type specific *Brd4* deletion in *Cx3cr1*-expressing cells and antibody-mediated IL1B neutralization.

## Conclusion

Organ function relies on a complex crosstalk between specialized cell types to maintain homeostasis and respond to injury and stress. Here, we leveraged both small-molecule BET bromodomain inhibition and cell-type specific *Brd4* genetic deletion in the context of heart failure to identify the stress-activated epigenomic mechanisms by which macrophages provide critical paracrine signals to activate cardiac fibroblasts in disease (**Fig. 5M**). Our work highlights how cell-compartment-specific perturbations of chromatin signaling can be combined with single-cell genomics to dissect the effects on individual cell states and elucidate critical signaling molecules governing cell-to-cell interactions in disease.

We show that deletion of *Brd4* in *Cx3Cr1*-positive macrophages is associated with improved cardiac function in a model of systolic heart failure. We observed not only a significant decrease in the proinflammatory transcriptional response in the monocyte-derived macrophages, but also an equally dramatic transcriptomic shift in fibroblasts, a population of cells which had not been genetically perturbed. In this context, the profibrotic response was abrogated, suggesting that in response to stress, *Cx3cr1*-expressing cells release potent paracrine signals, such as IL1B, that drive activation of tissue fibroblasts. Of note, although *Cx3cr1* is expressed in multiple immune cell subtypes, our transcriptome profiling suggests that monocyte-derived, *Ccr2*-positive macrophages also expressing *Cx3cr1* were the primary responders to BET inhibition and *Brd4* deletion. Other *Cx3cr1*-expressing populations, such as cardiac tissue-resident macrophages, might play a beneficial role in response to cardiac stress^36^. We found that IL1B was essential for fibroblast activation in heart failure pathogenesis, and that via a specific p65/RELA-dependent enhancer it controls the activation of MEOX1, a TF that we have recently identified as a critical regulator of the stress-activated fibroblast response^3^. While much of the study of fibroblast activation in disease has centered on the role of canonical profibrotic signaling pathways such as TGFB^31,32^, very little is known about how inflammatory signals can influence fibroblast cell states in disease. Our discoveries highlight how *cis*-regulatory elements such as the MEOX1 enhancer can function as hubs to process both profibrotic and proinflammatory signaling, providing a mechanistic understanding of how inflammation can modulate tissue fibrosis and remodeling.

Our data suggest that IL1B is a critical modulator of the stress-induced fibroblast states, and its antibody-mediated neutralization is sufficient to improve cardiac function in heart disease. Importantly, an exploratory analysis of a clinical trial evaluating systemic administration of an IL1B neutralizing antibody for treatment of ischemic cardiac disease^37^ identified a dose-dependent reduction in heart failure hospitalization and heart-failure related mortality^38^, spurring interest in the possibility of targeting IL1B signaling for the prevention or treatment of heart failure. As secreted factors such as IL1B can amplify the stress-response by acting on receptors expressed in non-immune cells, they represent exciting therapeutic targets that may dampen pathological cellular crosstalk and ameliorate organ dysfunction in disease.

While anti-inflammatory therapies targeting essential immune modulators have promising therapeutic applications, they are also associated with serious adverse events such as higher incidence of fatal infection and sepsis^37,39^. Here, by profiling *in vivo* BRD4 occupancy specifically in *Cx3cr1*-expressing macrophages under stress, and performing a series of CRISPR genomic deletions, we identified the exact stress-activated *cis*-elements that regulate *Il1b* activation in heart disease. This mechanistic refinement converging on specific regulatory elements offers key new insights that can catalyze the development of new therapeutic approaches to selectively inhibit proinflammatory states in defined cell compartments, which may be a preferable strategy for a wide variety of diseases associated with chronic inflammation and maladaptive tissue remodeling.

## Acknowledgments

We thank the Srivastava laboratory for critical discussions; Rudi Micheletti, Michael G. Rosenfeld, Karin Pelka, Gaia Andreoletti and Alexis J. Combes for helpful feedback; we thank Kathryn N. Ivey and Jin Yang at Tenaya Therapeutics for critical discussions and feedback; Jun Qi, Deyao Li and Logan Sigua for kindly providing the JQ1 molecule; we thank Jane Srivastava and all the Gladstone Flow Cytometry Core; we thank the Gladstone Animal Facility for support with mouse colony maintenance; and Ana Catarina Silva for help with figure editing and design.

## Author contributions

M.A., A.P. and D.S. conceived the study and interpreted the data. M.A. wrote the manuscript with inputs from A.P., S.M.H and D.S. Y.H. performed heart surgeries and echocardiography. M.A., A.P., Y.H., L.Y., C.Y.L., N.S, K.A., A.Z. and M.C. isolated cardiac cells for subsequent flow cytometry sorting and sc/snRNA- and ATAC-seq. M.A. and T.N. prepared chromium libraries. P.F.P and K.S.P developed computational methods for correlation and enhancer discovery analysis. M.A. analyzed scRNA-seq. T.N. and P.F.P analyzed scATACseq. M.A., A.P., N.S, B.G.T and R.J. performed and interpreted CUT&RUN. A.Pe. analyzed bulk RNA-seq and CUT&RUN data. J.G.T, K.L-C., R.J.V. and T.A.M performed and interpreted the studies on BMDMs, collagen-contractility and aSMA-immunofluorescence. L.Y. generated enhancer KO clones, and performed ChIP-seq and luciferase experiments. M.A., L.Y., N.S., W.F. and C.K-S.K. performed experiments on human iPS-CFs.

---

**Figure S1:**
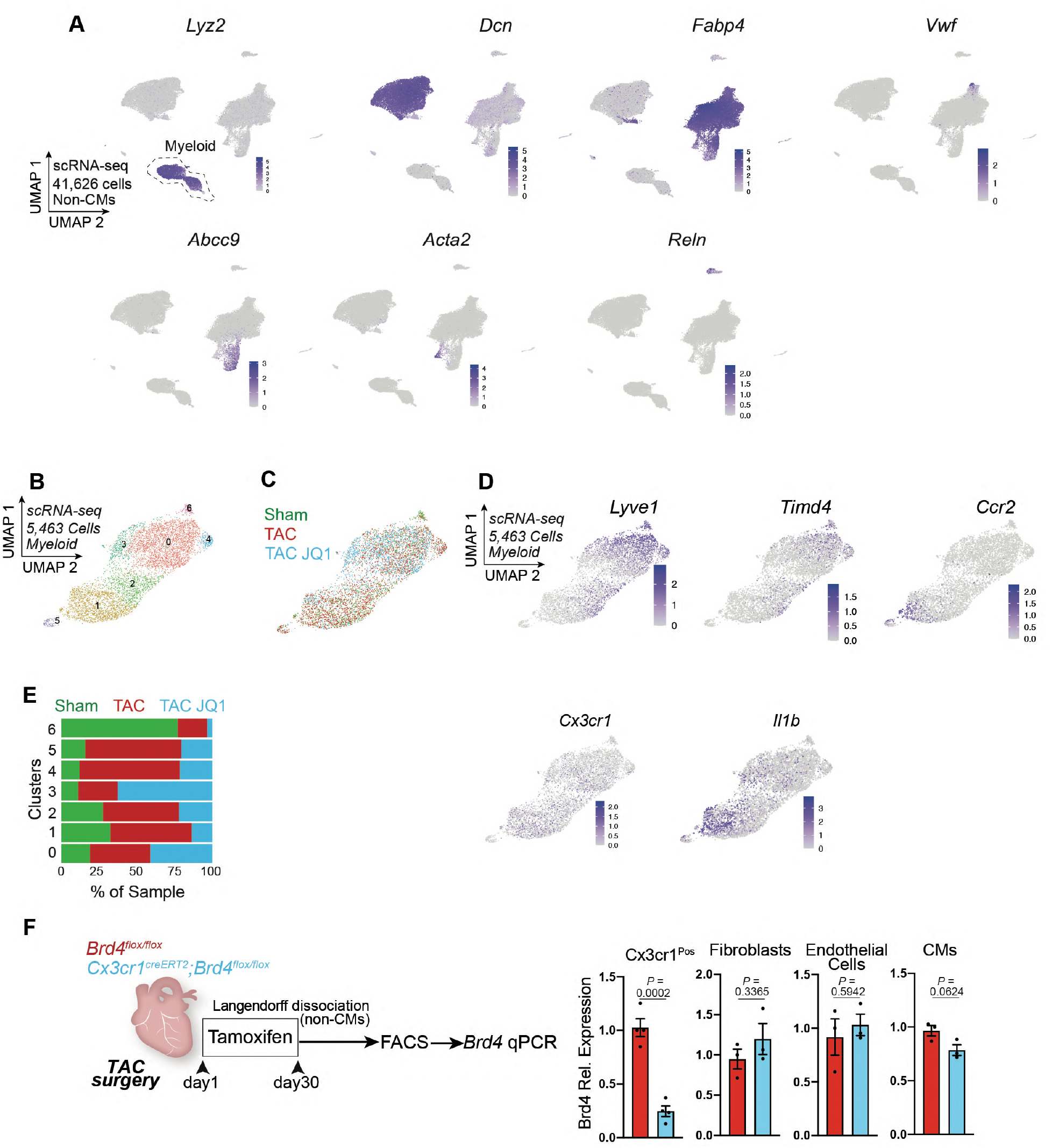
Single-cell RNA-seq in heart failure with BET inhibition identifies a highly dynamic Cx3cr1-expressing monocyte/macrophage subpopulation. **A.** Expression Feature Plots (scRNA-seq) of cell population markers in non-CM cells. **B,C**. UMAP plot (scRNA-seq) of myeloid cells colored by cluster (B) and sample identity (C). **D.** Expression Feature Plots (scRNA-seq) of Lyve1, Timd4, Ccr2, Cx3cr1 and Il-1b in myeloid cells. **E**. Sample distribution within clusters in myeloid cells. **F**. Schematic of experimental settings for conditional Brd4 deletion in Cx3cr1^Pos^ cells. Brd4 expression was measured by qPCR in sorted CX3CR1^Pos^, fibroblasts (mEF-SK4^Pos^, CD45^Neg^, CD31^Neg^), endothelial cells (mEF-SK4^Neg^, CD45^Neg^, CD31^Pos^) and in unsorted CMs. **F**, Data are mean ± s.e.m. Unpaired, two-tailed Student’s t-test.

**Figure S2:**
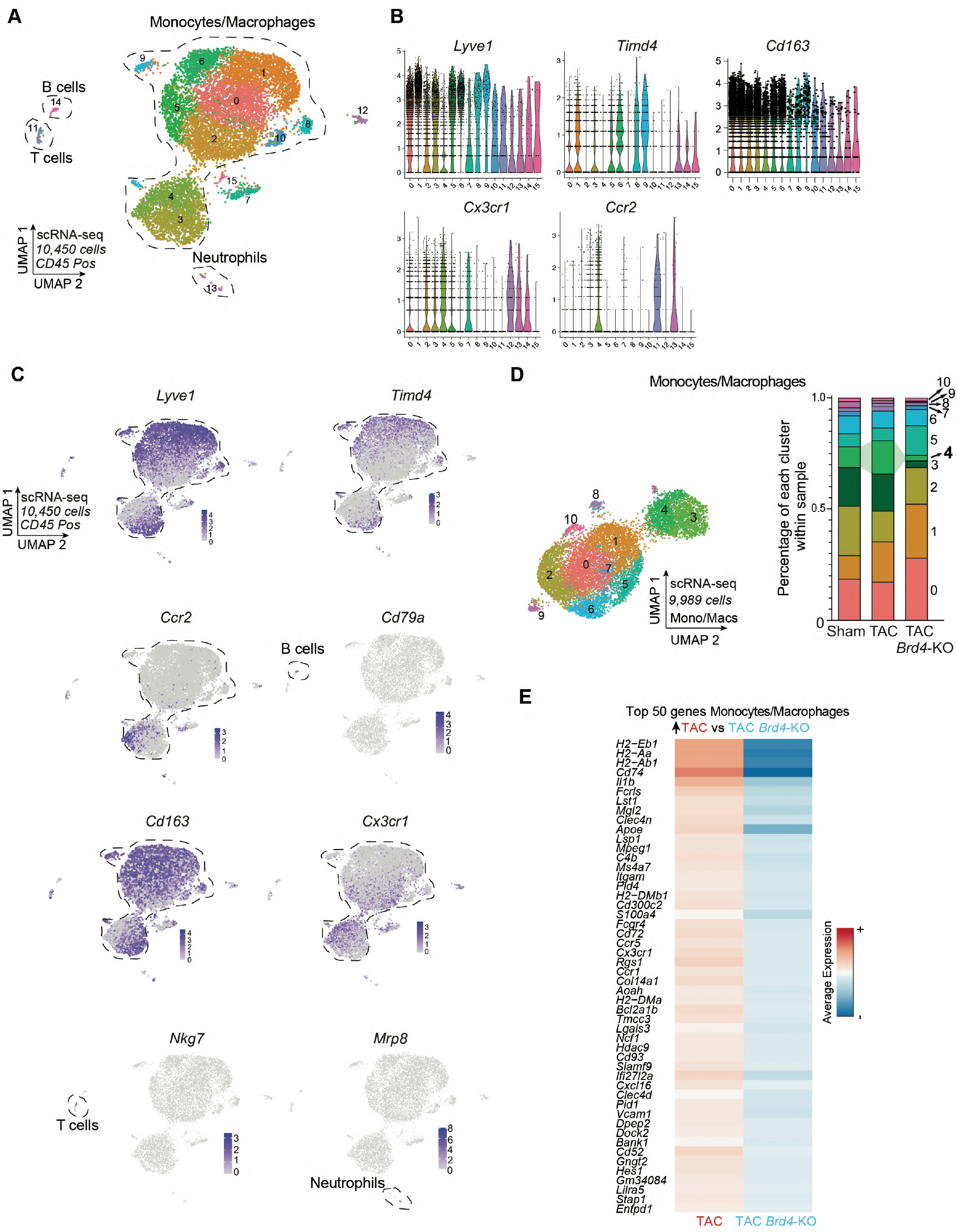
Identification of monocyte/macrophage population with single-cell RNA-seq. **A.** UMAP plot (scRNA-seq) of CD45^Pos^ cells colored by cluster identity. **B**. Expression Violin Plots of Lyve1, Timd4, Cd163, Cx3cr1 and Ccr2 in CD45^Pos^ cells. **C.** Expression Feature Plots (scRNA-seq) of Lyve1, Timd4, Ccr2, Cd79a, Cd163, Cx3cr1, Nkg7 and Mrp8 in CD45^Pos^ cells. **D.** UMAP plot (scRNA-seq) of monocytes/macrophages colored by cluster (left) and cluster composition within samples (right). Changes in cluster 4 are highlighted. **E.** Heatmap of expression (scRNA-seq) depicting the top 50 genes upregulated in TAC versus TAC Brd4-KO in the monocyte/macrophage population.

**Figure S3:**
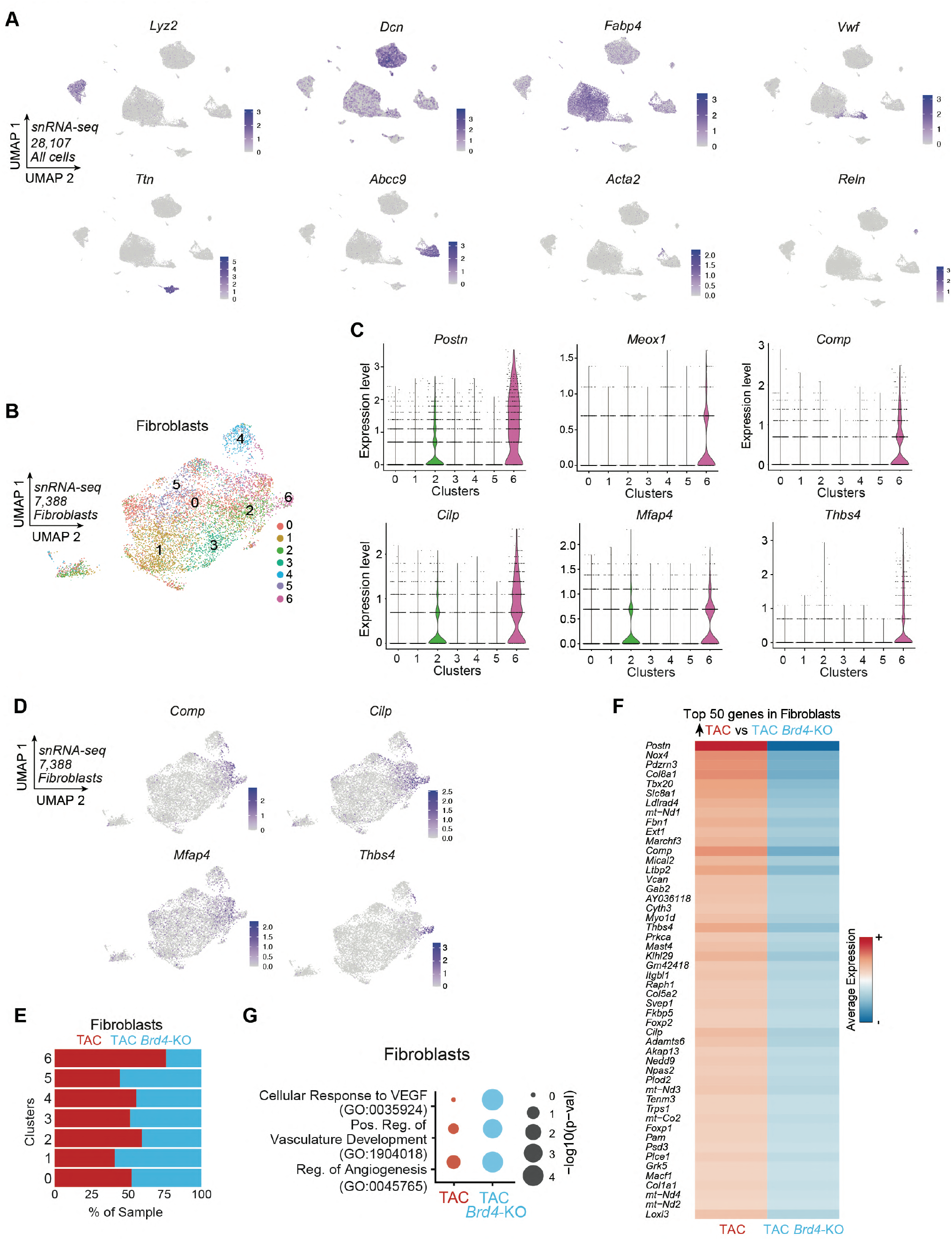
Decreased profibrotic signature in fibroblasts with Brd4 deletion in Cx3cr1-expressing monocytes/macrophages. **A.** Expression Feature Plots (snRNA-seq) of cell population markers in nuclei from cardiac tissue. **B**. UMAP plot (snRNA-seq) of fibroblasts colored by cluster identity. **C.** Expression Violin Plots (snRNA-seq) of Postn, Meox1, Comp, Cilp, Mfap4 and Thbs4 across clusters in fibroblasts. **D**. Expression Feature Plots (snRNA-seq) of Comp, Cilp, Mfap4 and Thbs4 in fibroblasts. **E**. snRNA-seq sample distribution within fibroblast clusters. **F**. Heatmap of expression (snRNA-seq) depicting the top 50 genes upregulated in TAC versus TAC Brd4-KO in fibroblasts. **G**. Dot Plot indicating significance (-log10(p–val)) for indicated GO terms in genes upregulated in TAC vs TAC Brd4-KO (red) or upregulated in TAC Brd4-KO vs TAC (blue) in fibroblasts.

**Figure S4:**
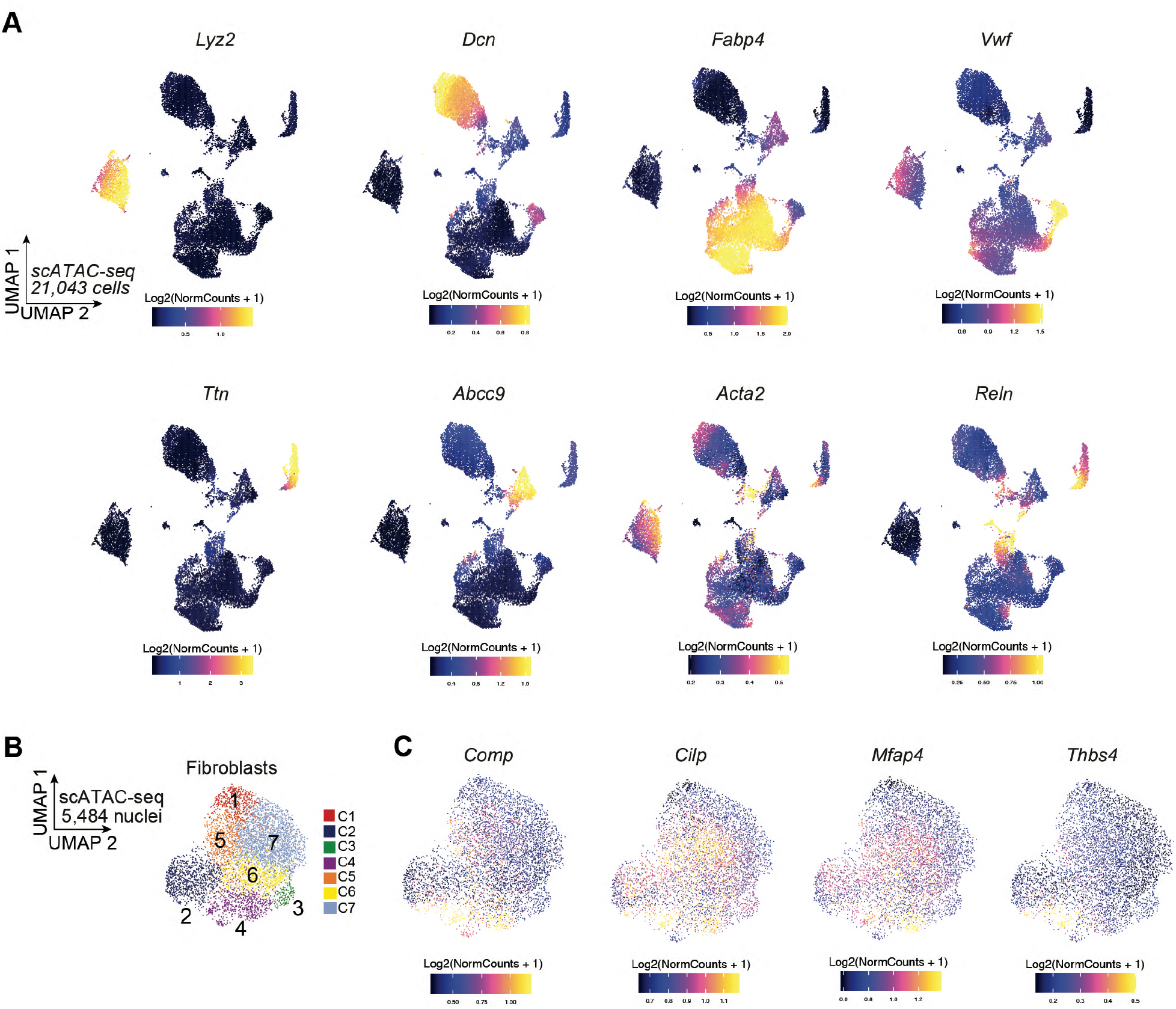
Changes in fibroblast chromatin accessibility with Brd4 deletion in Cx3cr1-expressing monocytes/macrophages. **A.** Chromatin accessibility gene score of cell population markers across nuclei from cardiac tissue. **B**. UMAP plot (scATAC-seq) of fibroblasts colored by cluster identity. **C.** Chromatin accessibility gene score of Comp, Cilp, Mfap4 and Thbs4 in fibroblasts.

**Figure S5:**
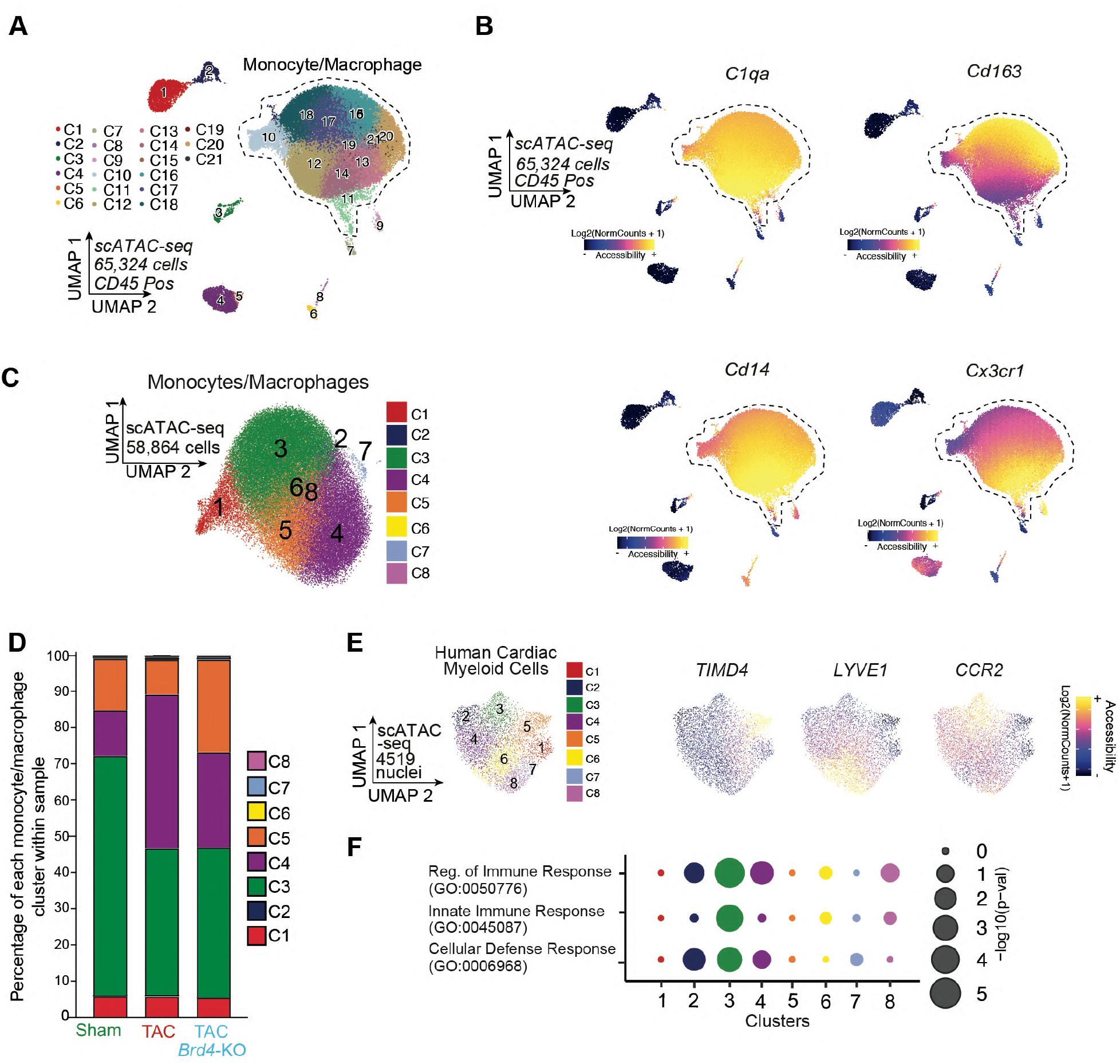
Identification of monocyte/macrophage population with single-cell ATAC-seq. **A.** UMAP plot (scATAC-seq) of CD45^Pos^ cells colored by cluster, the clusters encompassing the monocyte/macrophage populations are highlighted. **B**. Chromatin accessibility gene score of C1qa, Cd163, Cd14 and Cx3cr1 in CD45^Pos^ cells. **C.** UMAP plot (scATAC-seq) of monocytes/macrophages colored by cluster identity. **D**. Monocyte/macrophage cluster distribution within samples. **E**. Chromatin accessibility gene score of TIMD4, LYVE1 and CCR2 in human cardiac myeloid cells. **F**. Dot Plot indicating significance (-log10(p-val)) for indicated GO terms across human myeloid cell clusters.

**Figure S6:**
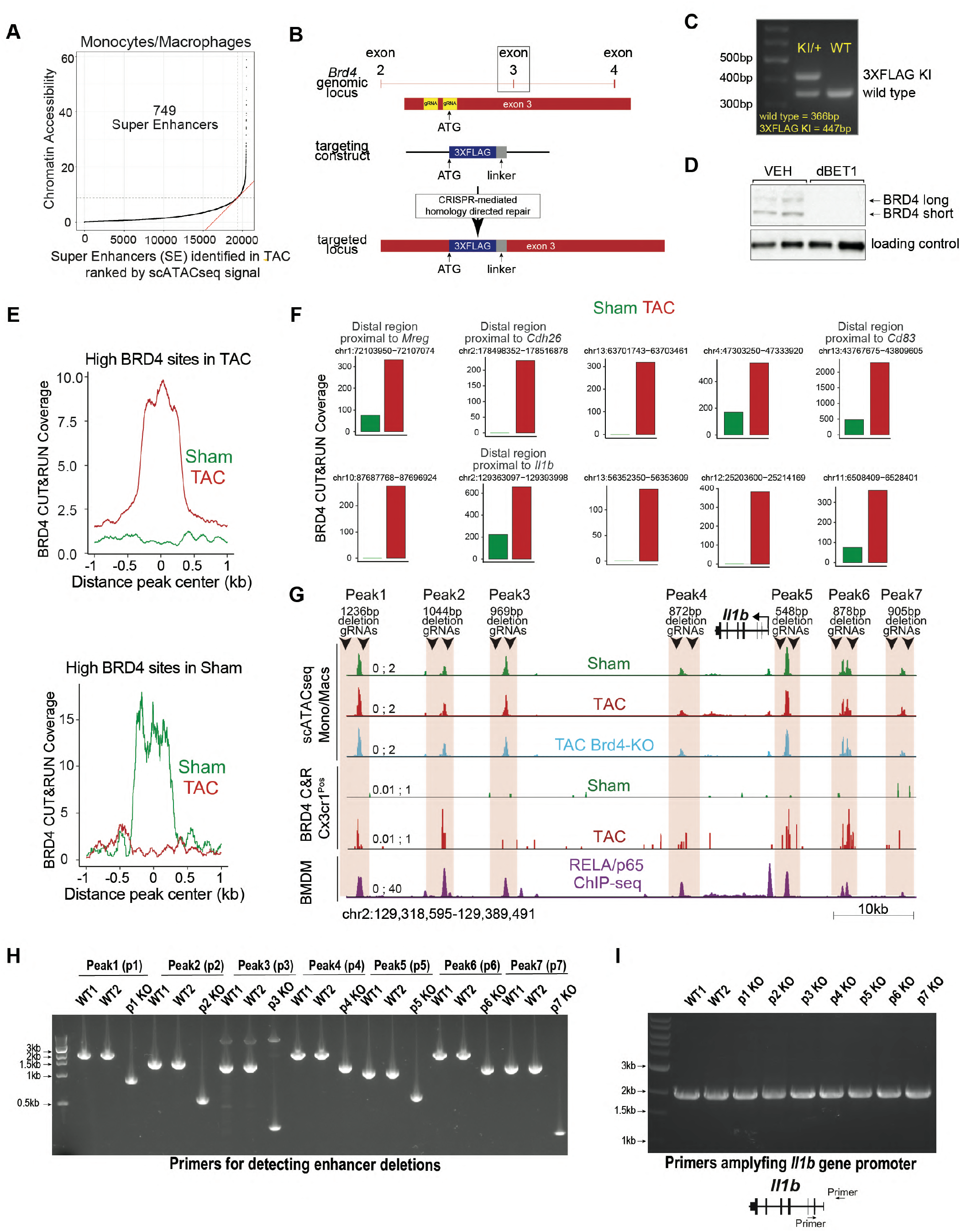
Chromatin accessibility and BRD4 occupancy in Cx3cr1-expressing cells identifies set of highly dynamic monocytes/macrophages distal elements in heart failure. **A**. Distribution of accessibility in monocytes/macrophages in the TAC state identifies a class of distal regions (superenhancers (SE)) for which the accessibility falls over the inflection point of the curve. **B**. Schematic of the targeting strategy to generate the Brd4 3xFlag mouse. **C.** Western blot showing WT and 3xFLAG knock-in bands in WT and Brd4^flag/+^ animals. **D**. Liver tissue western blot showing expression of endogenous long and short BRD4 isoforms treated with vehicle or with the small-molecule BET protein degrader dBET1^40^. **E**. Coverage from anti-FLAG CUT&RUN in Sham and TAC identifies regions enriched with BRD4 in stress (left) or at baseline (right). **F**. Coverage from anti-FLAG CUT&RUN in Sham and TAC in selected ten super enhancer regions. **G**. Schematic of the Il1b locus displaying the location of the gRNAs used to delete the 7 distal regions (Peak1 to Peak7). **H,I**. Agarose gel electrophoresis to assess distal peak deletions in Il1b locus (H) and unaffected region around Il1b promoter (I) (1813bp) in WT and KO clones.

**Figure S7:**
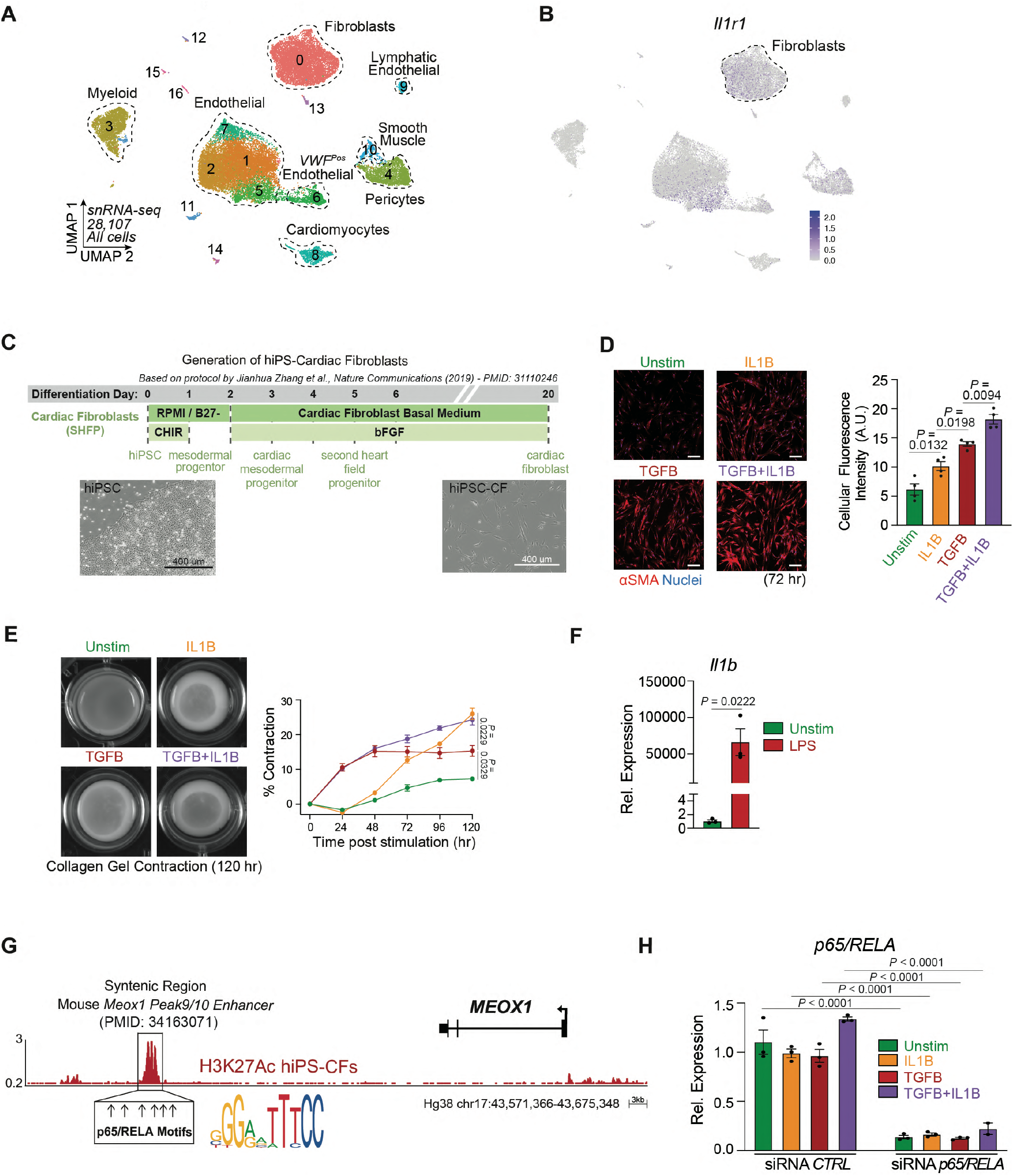
IL1B increases profibrotic response in human induced pluripotent cardiac fibroblasts. **A.** UMAP plot (snRNA-seq) of nuclei from cardiac tissue colored by cluster identity. **B**. Expression Feature Plot of Il1r1 in nuclei from cardiac tissue. Fibroblasts are highlighted. **C.** Protocol to generate human induced pluripotent cardiac fibroblasts (iPS-CFs). **D**. Immunofluorescence staining of αSMA (left) in Unstimulated iPS-CFs or treated with IL1B, TG-B or TGFB+IL1B. Nuclei are marked by Hoechst. Scale bars, 200 μm. Right, quantification of αSMA staining. **E**. Images (left) and quantification (right) of iPS-CFs seeded on compressible collagen gel matrices in unstimulated or with IL1B, TGFB or TGFB+IL1B treatments. **F**. Il1b expression by qPCR in Unstimulated and LPS treated Raw264-7 macrophages. **G**. Human MEOX1 locus showing H3K27Ac in unstimulated iPS-CFs. The syntenic region of the mouse Meox1 Peak9/10 regulatory element^3^ is highlighted and the six p65/RELA motifs within the region indicated. **H**. p65/RELA expression by qPCR in Unstimulated iPS-CFs or treated with IL1B, TGFB or TGFB+IL1B with control or p65/RELA-targeting siRNAs. **D**-**F**,**H** Data are mean ± s.e.m. One-way (D,H) and Two-way (E) ANOVA followed by Tukey post hoc test. Unpaired, two-tailed Student’s t-test (F).

**Figure S8:**
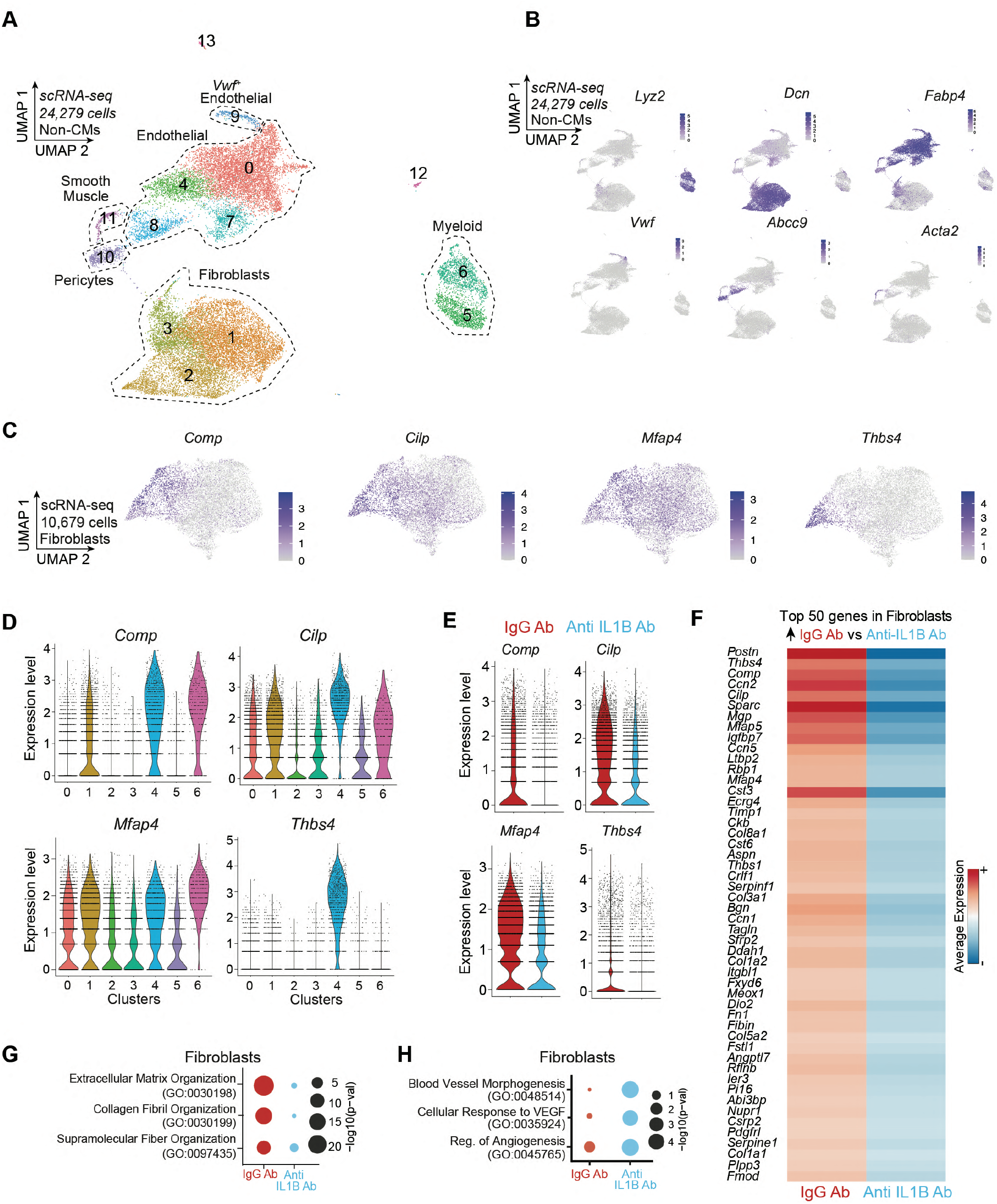
In vivo antibody-mediated IL1B neutralization decreases profibrotic transcriptional signature in fibroblasts. **A.** UMAP plot (scRNA-seq) of non-cardiomyocyte cells colored by cluster. **B** Expression Feature Plots (scRNA-seq) of cell population markers in non-cardiomyocyte cell population. **C**. Expression Feature Plots (scRNA-seq) of Comp, Cilp, Mfap4 and Thbs4 in fibroblasts. **D**. Expression Violin Plots (scRNA-seq) of Comp, Cilp, Mfap4 and Thbs4 across clusters in fibroblasts. **E**. Expression Violin Plots (scRNA-seq) of Comp, Cilp, Mfap4 and Thbs4 across samples in fibroblasts. **F**. Heatmap of expression (scRNA-seq) depicting the top 50 genes upregulated in TAC IgG Ab versus TAC Anti-IL1B Ab in fibroblasts. **G,H**. Dot Plot indicating significance (-log10(p-val)) for indicated GO terms in genes upregulated in IgG vs Anti-IL1B (red) or upregulated in Anti-IL1B vs IgG (blue) in fibroblasts.

## Methods

### Animal models

All protocols concerning animal use were approved by the Institutional Animal Care and Use Committees at the University of California, San Francisco and conducted in strict accordance with the National Institutes of Health Guide for the Care and Use of Laboratory Animals. Studies were conducted with age-matched male mice in a pure C57BL/6J background. Mice were housed in a temperature- and humidity-controlled facility with a 12-hour light/dark cycle. *Brd4^flox10^* and *Cx3cr1^CreERT219^* mice have been previously described. *Brd4^flag^* mice were produced by blastocyst injection of ribonucleoprotein complexes consisting of purified Cas9 protein (IDT), a guide RNA (crRNA targeting *Brd4* locus and universal 67mer tracrRNA; IDT), and a single-stranded oligonucleotide DNA template for homology directed repair (IDT) that led to the in-frame insertion of a 3X FLAG epitope tag along with a GGGGS flexible linker immediately downstream of the start codon in exon 3 of the endogenous *Brd4* locus (details of gRNAs and HDR are provided in the *Tables* section). Blastocysts were transferred to pseudopregnant females and pups were weaned at 4 weeks of age. Founder animals were screened for introduction of the 3X FLAG epitope tag by PCR amplification and confirmed by sequencing. Positive founders were outcrossed to wild type C57BL/6J animals and pups were screened for germline transmission of the *Brd4^flag^* allele by PCR genotyping with sequencing confirmation. Animals were outcrossed to C57BL/6J animals for >6 generations.

### Preparation of JQ1

JQ1 was synthesized and purified in the laboratory of Jun Qi (Dana Farber Cancer Institute), as previously published^4^. For *in vivo* experiments, a stock solution [50 mg/ml JQ1 in dimethyl sulfoxide (DMSO)] was diluted to a working concentration of 5 mg/ml in an aqueous carrier (10% hydroxypropyl b-cyclodextrin; Sigma C0926) using vigorous vortexing. Mice were injected at a dose of 50 mg/kg given intraperitoneally once daily. Vehicle control was an equal amount of DMSO dissolved in 10% hydroxypropyl b-cyclodextrin carrier solution. All solutions were prepared and administered using sterile technique. For *in vitro* experiments, JQ1 was dissolved in DMSO and administered to cells at 500nM final concentration using an equal volume of DMSO as control.

### Mouse model of transverse aortic constriction and echocardiography

All mice were male and 10-12 weeks of age. Wild type C57BL/6J mice were obtained from The Jackson Laboratory (Stock No: 000664). Mice were placed on a temperature-controlled small-animal surgical table to help maintain body temperature (37°C) during surgery. Mice were anesthetized with isoflurane, mechanically ventilated (Harvard Apparatus), and subjected to thoracotomy. For TAC surgery, the aortic arch was constricted between the left common carotid and the brachiocephalic arteries using a 7-0 silk suture and a 25-gauge needle, as previously described^5^. For sham surgeries, thoracotomy was performed as above, and the aorta was surgically exposed without any further intervention. For echocardiography, mice were anesthetized with 1% inhalational isoflurane and imaged using the Vevo 3100 High Resolution Imaging System (FujiFilm VisualSonics Inc.) and the MX550S probe. Measurements were obtained from M-mode sampling and integrated electrocardiogram-gated kilohertz visualization (EKV) images taken in the left ventricle (LV) short axis at the mid-papillary level as previously described^5^. LV ejection fraction measurement was obtained from high-resolution two-dimensional measurements at end-diastole and end-systole as previously described^5^. All echocardiography analyses were performed blinded (mice were assigned an alphanumeric code).

### Flow cytometry analysis and cell sorting

Non-CMs purified from Langendorff perfused hearts were suspended in 500ul of FACS buffer (5% FBS, 0.01% NaN3 in 1X PBS) and blocked with Fc receptor antibody specific for FcγR III/II (Biolegend TruStain FcX PLUS anti-mouse CD16/32; 1:50 dilution) for 15 minutes, then stained with indicated antibodies for 30-40 minutes. After staining, cells were washed three times with FACS buffer and resuspended in 400ul of 5% FBS in 1X PBS. Stained cells were subjected to flow cytometry using a BD FACSAria II cell sorter. Antibodies used in this study are mCD45-PB (Biolengend; 1:100), mCx3cr1-APC (Biolegend; 1:100), CD31-PE-Cy7 (Invitrogen; 1:50), and mEFSK4-APC (Miltenyi Biotec; 1:40).

### Langendorff perfusion and cell/nuclei isolation for subsequent single-cell RNA and ATAC sequencing

Cell isolation from mouse hearts were performed using two methodologies: #1) as previously described with minor modifications^41^ or #2) using a variation of the PAN-INTACT protocol^42^. For #1, after a proper level of anesthesia was induced, a thoracotomy was performed and the mouse heart was isolated, cannulated, and perfused with perfusion buffer (120.4 mM NaCl, 14.7 mM KCl, 0.6 mM KH2PO4, 0.6 mM Na2HPO4, 1.2 mM MgSO4, 10 mM Na-HEPES, 4.6 mM NaHCO3, 30 mM taurine, 10 mM 2,3-butanedione monoxime, and 5.5 mM glucose, pH 7.0) in a Langendorff perfusion system (Radnoti 120108EZ) for 5-10 minutes at 37°C. The cannulated heart was then digested with digestion buffer (perfusion buffer with 300 units/mL collagenase II (Worthington Biochemical) and 50 μM CaCl2) for approximately 10 min at 37°C. At the end of digestion, the atria and great vessels were removed and the ventricular tissue was transferred and gently teased into small pieces in stop buffer (perfusion buffer with 10% fetal bovine serum) at 37°C. After gently pipetting, the cell suspension was passed through a 250 μM strainer in a 50ml conical centrifuge tube and centrifuged at 30x*g* for 3 minutes at room temperature (RT). The supernatant, which contained most of the non-cardiomyocytes (non-CMs), was divided from the pellet (which contained the CM fraction). The non-CM fraction was centrifuged again at 30x*g* for 3 minutes at RT and the supernatant retained. The supernatant was filtered with a cell strainer (70 μm) and centrifuged at 400x*g* for 3 minutes at RT for eliminating debris. The non-CM pellet was resuspended in 1mL cold PBS 0.5% BSA. Cells were counted with trypan blue using a hemocytometer and 15k cells were used for subsequent 10X Genomics Chromium single-cell RNAseq preparation. For single cell ATAC, 500k isolated and purified non-CMs were resuspended in 100uL lysis buffer (Tris-HCl 10mM pH 7.4, NaCl 10mM, MgCl2 3mM, Tween-20 0.1%, P40 0.1%, Digitonin 0.01%, BSA 1% in nuclease-free water), pipetted 10 times, and kept on ice for 5 minutes. Nuclei were washed with 1mL 1X PBS with 1% BSA and centrifuged for 5 minutes at 4 °C. The nuclei pellet was resuspended in 1mL 1X PBS with 1% BSA and filtered with a 10uM strainer (pluriSelect #43-50010-00). Nuclei were counted after DAPI staining using a hemocytometer and 10k non-CM nuclei were used for subsequent 10X Genomics Chromium single-cell ATAC-seq preparation. The CM fraction underwent a second round of centrifugation at 30x*g* for 3 minutes at RT in stop buffer and the supernatant was discarded. The CM pellet was finally centrifuged at 400x*g* for 3 minutes in stop buffer at RT and the supernatant was discarded. For #2, after the mouse heart was isolated it was rinsed in a sterile 10cm petri dish containing 1X PBS on ice to remove excess blood and transferred to a 10cm petri dish containing 2ml of ice cold lysis buffer (320 mM sucrose, 5 mM CaCl2, 3 mM magnesium acetate, 2 mM EDTA, 0.5 mM EGTA, 10 mM Tris-HCl pH 8.0, 0.20% triton X-100) and minced into very small pieces. The minced heart tissue in 2ml of lysis buffer was transferred to a “Type A” dounce homogenizer and subjected to 50 strokes. This solution was subsequently transferred to a “Type B” dounce homogenizer and subjected to 3 strokes. The solution was serially passed through 100 μM and 70 μM strainers and then centrifuged at 1,000x*g* at 4 °C for 8 minutes. The nuclear pellet was resuspended in nuclei resuspension buffer (430 mM sucrose, 70 mM KCl, 2 mM MgCl2, 10 mM Tris-HCl ph 7.2, EGTA 5 mM, 10% glycerol). Nuclei were counted after DAPI staining using a hemocytometer and 15,000 nuclei were used for subsequent 10X Genomics Chromium single-nuclei RNAseq and 10,000 nuclei were used for ATAC-seq.

### sc/snRNA-seq and scATAC-seq library preparation

For sc/snRNA-seq library preparation, cells or nuclei were loaded onto the 10X Genomics Chromium instrument according to manufacturer’s protocols (Chip G, PN-1000120). All experiments were conducted with version 3.1 NEXT GEM reagents (PN-1000121), using 9 cycles of cDNA amplification for the GEM kit (PN-1000123) and 9 cycles of library amplification for library kit (PN-1000157). Each sample was indexed with a unique sample primer (Single Index Kit T Set A, PN-000213). For scATAC-seq library preparation, nuclei were first transposed with Tn5 enzyme (PN-2000138) and then loaded onto the Chromium instrument (Chip H, PN-1000161) using the version 1.1 NEXT GEM reagents (PN-1000175). Library preparation proceeded with 12 cycles each of GEM barcoding and library amplification. Each sample was indexed with a unique sample primer (Single Index Plate N Set A, PN-3000427). For both scRNA-seq and scATAC-seq, samples were pooled separately into respective libraries and sequenced on an Illumina NextSeq 500 (Illumina, software 4.0.2) and/or a lane of a Novaseq S4 (Illumina, software v1.5), according to manufacturer’s guidelines for sc/snRNA-seq and scATAC-seq, respectively.

### sc/snRNA-seq data preparation and analysis

The Cell Ranger pipeline was used for processing all samples post-sequencing (10X Genomics, version 3.1). Samples were demultiplexed and fastq files were generated using cellranger mkfastq. All samples were then individually aligned to the mouse reference genome mm10 using cellranger count with the intron = true flag to be consistent with single nuclei RNA-seq analysis for which intronic reads are also kept. To account for varying sequencing read depth, we further ran cellranger aggr to normalize all samples to mapped read depth of the least sequenced sample. Read-normalized aggregated values from cellranger aggr were used as initial input using the R package Seurat (v4.0.1) using the functions Read10X and CreateSeuratObject with standard variables. Each sample used in aggregation was identified by condition and replicate and assigned a unique name in metadata as “gem.group”. Quality control filtering included removal of outliers due to the number of features/genes (nFeature_RNA > 2000 & nFeature_RNA < 7500), UMI counts (<80,000) and mitochondrial percentage (<15%). SCT normalization was then performed. PCA analysis and batch correction using Harmony was then performed using split.by = “gem.group”. Clustering was then run using the functions RunUMAP, FindNeighbors, and FindClusters and the output UMAP graphs were generated by DimPlot. Marker genes were identified by the function FindAllMarkers with standard settings. After initial processing, iterative rounds of filtering poor quality clusters and re-running clustering workflows. Differential gene expression was performed using the Wilcoxon test between two groups with the function FindMarkers. Gene ontology (GO) analysis was run using genes identified with either FindAllMarkers or FindMarkers using Enrichr ^43^.

### Correlating cardiac function with gene expression

To identify genes correlated with changes in cardiac function across conditions, we designed a regression model to link cell type specific changes between reads in scRNA-seq data to the measured ejection fraction of the left ventricle by echocardiographic assessment. For each gene, we fit a linear model where the output variable is the number of reads in that gene for each cell of the particular cell type of interest and the input is the ejection fraction matched to the same condition that the cells are taken from, using a Poisson distribution to model the counts. We sampled down to 1,000 cells in each cell type and condition, resampling when there were too few cells. A score is assigned to each gene in each cell type using the correlation coefficient of the fitted regression on the ejection fraction. We then normalized the scores for each cell type across genes using the interquartile range, and used the variance of the normalized scores to compute *p*-values. We use a threshold of +/- 5 (*p*-value < 1e^-6^) to consider a gene to be significantly positively or negatively correlated with cardiac function in a given cell type.

### scATAC-seq data preparation and analysis

Raw sequencing data were preprocessed with the Cell Ranger ATAC v.2.0 pipeline (10X Genomics). The ArchR v1.0.1 R package^44^ was used for subsequent analyses according to the ArchR web tutorials. The TSS enrichment score and the number of fragments per nuclei were used to eliminate low quality nuclei from downstream analyses (for *Cx3cr1^creERT2^;Brd4^flox/flox^* whole heart ATAC-seq data (Fig. 3), minTSS=16, minFrags=3163, and maxFrags=1e+6; For *Cx3cr1^creERT2^;Brd4^flox/flox^* CD45^Pos^ ATAC-seq data (Fig. 4), minTSS=15, minFrags=1584, and maxFrags=1e+6), and nuclei doublets were removed using the addDoubletScores and filterDoublets functions, resulting in the ArchRProject to be analyzed. After genome-wide tiling with iterative Latent Semantic Indexing, dimensionality reduction was performed using the addIterativeLSI function. Batch correction among each sample was performed using the addHarmony function. Clustering and the 2D embedded visualization in UMAP space were performed using the addClusters, addUMAP, and plotEmbedding functions, respectively. For *Cx3cr1^creERT2^;Brd4^flox/flox^* CD45^Pos^ ATAC-seq, some clusters prominently enriched with only one sample from Cre positive animals and those nuclei showed lower TSS and lower nFragments than those of nuclei from other clusters. Thus, those clusters were excluded as low quality data with the remainder of the nuclei processed again in the same manner. Gene scores (GS), which were calculated by the accessibility of promoter and gene body regions of each gene and can be treated as a proxy of expression levels of a corresponding gene, were extracted to identify the cluster features using the getMarkerFeatures with useMatrix=“GeneScoreMatrix”. For peak calling per cluster, the addGroupCoverages and the addReproduciblePeakSet functions with peakMethod=“Macs2” were used. To identify the cluster specific feature peaks, the getMarkerFeature functions with useMatrix=“PeakMatrix” was used. Differentially accessible region (DAR) analysis between clusters was performed using the getMarkerFeatures function by setting clusters for comparison using the useGroups and bgdGroups functions. The statistically significant DARs were defined with FDR<0.1 and Log2 FC>1. Motif enrichment analyses on the detected DARs were performed using the peakAnnoEnrichment function. To identify the enriched transcription factor binding site motifs in the detected specific feature peaks found in our ATAC-seq analyses, the findMotifsGenome.pl function from HOMER^45^ was used with “-size given” and “-mask” options.

### Downstream processing of ATAC-seq data

For downstream analysis of ATAC-seq data we use ArchR generated cell-by-bin matrices. We normalize this matrix per cell using each cell’s total reads in transcription start sites (TSS), generating TSS normalized read counts for each cell in each bin. To generate coverage tracks, we use the mean TSS normalized accessibility across cells for the given cell type and condition at each bin. To compare the accessibility of a genetic region across conditions we use the mean of TSS normalized accessibility of cells across bins overlapping the region with a confidence interval computed on the deviation in accessibility across cells. To fit a curve to the change in accessibility over conditions, we fit a spline with two degrees of freedom to the TSS normalized accessibility of cells in each condition. To compute the co-accessibility of a peak with a promoter we use the Jaccard similarity of the binarized version of the ArchR cell-by-bin matrix, counting any bin overlapping the promoter and any bin overlapping the peak of interest.

### Identifying stress sensitive distal elements

To identify distal elements in the monocyte/macrophage population that are sensitive to stress and *Cx3cr1*-specific *Brd4* deletion we first use a modified version of the ROSE^25^ algorithm on all TAC distal peaks to stitch together peaks using a standard 12.5kb gap. However, in our version, we allow stitching to span gene coding regions but do not consider peaks in those regions. We score each stitched region using the sum of TSS normalized accessibility across the region, omitting reads in gene coding regions. We then use the standard ROSE algorithm to select large (or “super”) enhancers based on setting a threshold using the tangent line to the elbow plot of rank against score. This results in 749 large enhancers. We then score these for significance at the transition between Sham and TAC and between TAC and *Cx3cr1*-specific *Brd4* deletion as follows: at each transition we compute the TSS normalized number of reads for each cell in that region. We then use a one-tailed Wilcoxon rank-sum test in each direction to compute the significance of the change in accessibility across cells between each pair of conditions. We called two sets of dynamic SEs, ones that increase significantly (Wilcoxon *p*-value < 1e^-5^ in all cases) with TAC and decrease with *Cx3cr1*-specific *Brd4* deletion, and ones that decrease with TAC and increase with the deletion.

### Assessment of BRD4 chromatin occupancy by CUT&RUN in *Cx3cr1^Pos^* sorted cells

Whole hearts were obtained from *Brd4^flag/flag^* animals that underwent Sham or TAC surgeries and were subjected to Langendorff perfusion to obtain suspensions of cardiac cells. The non-CM population was purified by serial differential centrifugation and size-exclusion filtering steps (see relevant section on the Methods) followed by CX3CR1 antibody staining and sorting by flow cytometry (see dedicated section on flow cytometry for details). CX3CR1^Pos^ cells were subjected to CUT&RUN using the CUTANA^™^ ChIC/CUT&RUN kit (Epicypher) using an anti-FLAG antibody (Sigma M2 #1804) following manufacturer instructions with included positive and negative controls. Purified fragmented DNA was quantified using a Qubit 2.0 fluorometer (ThermoFisher Scientific) using the dsDNA HS Assay Kit. Paired-end Illumina sequencing libraries were prepared from CUT&RUN enriched DNA using the NEBNext Ultra™ II DNA Library Prep Kit (New England Biolabs) and sequenced on a NextSeq 500 (Illumina). The nf-core/cutandrun (v2.4.2 https://doi.org/10.5281/zenodo.5653535) pipelines were used to perform primary analysis within the Nextflow bioinformatics workflow manager (v21.10.6) in conjunction with Singularity (v2.6.0) using the following command: ‘nextflow run nf-core/cutandrun --genome GRCm38 --input design_matrix.csv-profile singularity’. The pipelines perform adapter trimming, read alignment, and filtering, normalized coverage track generation, peak calling and annotation relative to gene features, consensus peak set creation, and quality control and version reporting. All data were processed relative to the mouse (GCRm38 - mm10) annotation.

### Calculating BRD4 enrichment

For each SE (super enhancer) we calculated whether it had an increase or decrease in BRD4 binding. In order to score based on creation of new binding rather than just increase in overall binding, we compute the log-fold change (with a pseudocount of 1) in the total sum of the widths of detected BRD4 peaks in the two conditions. In order to conduct genome-wide scoring of changes in BRD4 binding we used the “csaw” package for R to bin the entire genome into 500bp windows and fit a model using the standard procedure and default parameters. This gave us a normalized *p*-value and log-fold change in binding for every 500bp window.

### ChIPseq experiment and analysis

For chromatin immuno-precipation (ChIP) sequencing experiments, 10^6^ human induced pluripotent stem cell derived cardiac fibroblasts (iPSC-FBs) were pelleted in unstimulated condition and resuspended in 10ml FibroGRO^™^ Basal Medium, cross-linked in 1% formalin solution (Thermo Fisher Scientific) by rocking in room temperature for 10 minutes and finally glycine (final concentration 0.125M) was added to quench the cross-link for 5 minutes. Samples were centrifuged at 1000 rcf for 5 minutes at 4C. Cells were washed twice with 10 ml of cold 1x PBS supplemented with proteinase inhibitors and phosphatase inhibitors (Roche # 4693132001) and the pellets were snap frozen in liquid nitrogen. All samples were stored at −80C until use. When ready, cell pellets were incubated in cell lysis buffer (20 mM Tris-HCl, pH 8, 85 mM KCl, 0.5% NP-40 and protease inhibitors) for 10 min on a rotator at 4C. Nuclei were isolated by centrifugation (2,500 x g, 5 min, 4C), resuspended in 1mL nuclear lysis buffer (50 mM Tris-HCl, pH 8, 10 mM EDTA, pH 8, 1% SDS, protease inhibitors) and incubated on a rotator for 30 min at 4C. Chromatin was sheared using a Covaris S2 sonicator (Covaris Inc) for 20 min (60 s cycles, 20% duty cycle, 200 cycles/burst, intensity = 5) for preparing DNA in the 200–700 base-pair range. Chromatin was diluted 3-fold in ChIP dilution buffer (0.01% SDS, 1.1% Triton X-100, 1.2mMEDTA, 16.7mMTris-HCl, pH 8, 167 mM NaCl, protease inhibitors) for a total volume of 3ml. 30uL of chromatin (representing 1%) was taken as input DNA. High-salt buffer (250mM Tris-HCl, pH 7.5, 32.5 mM EDTA, pH 8, 1.25M NaCl) and Proteinase K (New England Biolabs Inc (NEB)) were added and crosslinks were reversed overnight at 65C. Inputs were treated with RNase A, and DNA was purified with AMPure XP beads (Beckman Coulter cat #A63881). DNA concentration was measured with Qubit (ThermoFisher) to work out the original concentration of the sonicated chromatin. 15ug of chromatin was incubated with 4ug of anti-p65/RELA antibody (Bethyl Laboratories #A301-824A) and 3ug of chromatin was incubated with 2ul of anti-H3K27ac (Abcam #4729) for H3K27ac ChIP. All incubations were overnight at 4C under rotation. Antibody-protein complexes were immunoprecipitated using Pierce Protein A/G magnetic beads at 4C for 2 h under rotation. Beads were washed five times (2-min/wash under rotation) with cold RIPA buffer (50 mM HEPES-KOH, pH 7.5, 500 mM LiCl, 1 mM EDTA, 1% NP-40, 0.7% Na-deoxycholate), followed by one wash in cold final wash buffer (1xTE, 50 mM NaCl).

Immunoprecipitated chromatin was eluted at 65C with agitation for 30 min in the elution buffer (50mMTris-HCl pH 8.0, 10mMEDTA, 1% SDS). The immunoprecipitated chromatin was then reverse-crosslinked and purified as described above for the inputs. For subsequent sequencing, fragmented ChIP and input DNA were end-repaired, 5’-phosphorylated and dA-tailed with NEBNext Ultra II DNA Library Prep Kit for Illumina (NEB, E7645). Samples were ligated to adaptor oligos for multiplex sequencing (NEB, E7335), PCR amplified, and sequenced on a NextSeq 500 (Illumina, software 4.0.2) at the Gladstone Institutes.

The nf-core/ChIP-seq pipeline (v1.2.2; https://doi.org/10.5281/zenodo.4711243) was used to perform primary analysis of the samples within the Nextflow bioinformatics workflow manager (v21.10.6) in conjunction with Singularity (v2.6.0) using the following command: ‘nextflow run nf-core/chipseq --max_memory 80.GB --single_end --narrow_peak --skipBiotypeQC --skipTrimming --genome GRCh38/GRCm38 --input design_matrix.csv-profile singularity’. The pipeline performs adapter trimming, read alignment, and filtering, normalized coverage track generation, peak calling and annotation relative to gene features, consensus peak set creation, differential binding analysis, and quality control and version reporting. All data were processed relative to the human (GCRh38 - hg38) annotation.

### Lipopolysaccharide (LPS) stimulation of Raw 264-7 macrophages

The Raw 264-7 mouse macrophage cell line was obtained from the American Type Culture Collection (ATCC) and cultured in high-glucose Dulbecco’s Modified Eagle’s Medium (DMEM) containing 10% fetal bovine serum (FBS; Gibco), 100U/ml penicillin (Gibco), 100ug/ml streptomycin (Gibco), 1X non-essential amino acid solution (Gibco), 1mM sodium pyruvate (Gibco) at 37°C in a humidified incubator with 5% CO2. Approximately 2×10^5^ cells were seeded on 6-well plates in the late afternoon and cultured overnight. The following day, the culture medium was removed and replaced with fresh medium containing 0.1ng/ml, 1ng/ml, or 10ng/ml LPS (Sigma Aldrich). After 3, 6, or 24 hours following stimulation, cells were washed with 1X PBS and lysed in 1ml of Trizol LS reagent (Invitrogen) and processed for total RNA extraction.

### Generation of *Il1b* enhancer peak deletions in Raw 264-7 macrophages

We used CRISPR/Cas9-mediated genome editing to generate individual Raw 264-7 cell lines containing deletions of each of the 7 peaks identified from our scATAC-seq analysis. Chemically modified synthetic single guide RNAs (sgRNAs) flanking the region to be deleted (Synthego) were reconstituted in nuclease-free water at 100 uM. The paired ribonucleoprotein (RNP) complexes for each peak to be deleted were prepared individually by mixing 120 pmol of sgRNA with 20 pmol of Cas9 protein (QB3 MacroLab, University of California, Berkeley) in 5 ul of P3 primary cell nucleofection buffer (Lonza) and incubated at room temperature for 15 minutes. Prior to nucleofection, the paired RNPs were combined and nucleofected into 4×10^5^ Raw 264-7 cells suspended in 10 ul of nucleofection buffer (Lonza) using the program DS-136 in a 4D Nucleofector X Unit (Lonza). After nucleofection, 100 ul of warm culture medium was added to the cuvette and incubated at 37°C for 15 minutes. Nucleofected cells were equally divided to 4 wells of 96-well plate and cultured for 48 hours at 37°C. Cells from each of these 4 wells were dissociated and divided into 2 portions–one portion was used to collect genomic DNA using QuickExtract DNA Extraction solution (Lucigen) while 500 cells from the other portion was suspended in 25 ml of culture medium and 100 ul was seeded into each well of two 96-well plates for further culture. The extracted genomic DNA was subject to PCR to detect deletion of the desired regions and the products were visualized using agarose gel electrophoresis. The well with the highest density product was chosen for subsequent screening from the corresponding 96-well plates. In the 96-well plates, those wells observed to have single cells on the day after seeding were marked and maintained in culture for 7-10 days with medium changes every 2 days. When cells were 50% confluent, they were split into two wells in separate plates at different densities. The plate seeded at the higher density was harvested on the day after seeding and processed for genomic DNA extraction and PCR genotyping for detection of the targeted deletion. PCR products with smaller sizes than wild type band were verified by Sanger sequencing (Quintara Bio). Upon confirmation of the appropriate enhancer deletion, the corresponding lower density wells were expanded. The sequences of the sgRNAs used for individual peak deletions and the primers used to detect these deletions by PCR are outlined in the relevant “Tables” section.

### Analysis of publicly available human scATAC-seq data

After the human scATAC-seq data^28^ was processed and clustered by ArchR, we called marker genes for each cluster using getMarkerFeatures with Gene Scores with the wilcoxon test for each cluster at FDR<0.15 and a minimum log2-fold change in score of 0.5. To generate coverage tracks for the *IL1B* locus we selected all cells in Cluster 3 (n=632, the cluster with high accessibility for both *CX3CR1* and *IL1B* genes) and then calculated the mean TSS normalized coverage (as described previously) for cells derived from control patients and myocardial infarction (MI) patients. We plotted the coverage for the *IL1B* human locus (Hg38 chr2:112,798,605-112,843,614) obtained by lifting over the *Il1b* mouse locus (Mm10 chr2:129,318,595-129,389,491). To control for the large variance that can result from deriving coverage from a small number of cells, we only considered signal in regions in which at least two different samples had at least one read, and set a minimum threshold of 4e^-6^ for detectability of coverage

### Human induced pluripotent stem cell derived cardiac fibroblasts (iPSC-FBs)

Human induced pluripotent stem cell derived cardiac fibroblasts (iPSC-FBs) were cultured in FibroGRO Complete Medium (SCMF001; Millipore Sigma) with 2% FBS at 37°C in a humidified incubator with 5% CO2. When stimulating stress in iPSC-CFs, the cells were cultured with 0.5% FBS in FibroGRO Complete Medium without supplements, with addition of either TGFB (5ng/ml; Peprotech), IL1B (10ng/ml; Sigma Aldrich), or TGFB+IL1B (10ng/ml and 5ng/ml, respectively). For luciferase experiments, HeLa cells were obtained from ATCC and cultured in high-glucose Dulbecco’s Modified Eagle’s Medium (DMEM) containing 10% FBS (Gibco), 100U/ml penicillin (Gibco), and 100ug/ml streptomycin (Gibco) at 37°C in a humidified incubator with 5% CO2.

### RNA extraction, RT-PCR, and real-time PCR analysis

Total RNA was extracted from human iPSC-FBs and mouse Raw 264-7 cells using the Direct-Zol RNA kit (Zymo Research) according to manufacturer instructions. 500 ng of RNA was converted to cDNA using iScript cDNA synthesis kit (BioRad). For Taqman real-time PCR analysis, 1/50 cDNA was used for quantitative PCR with Taqman Universal PCR master mix (Life Technologies) and the QuantStudio 5 Real-Time PCR System (ThermoFisher Scientific). The Taqman probes and SYBR primers are outlined in the relevant “Tables” section.

### Luciferase reporter assays

The human *MEOX1* enhancer syntenic to the mouse *Meox1* peak9/10 enhancer^3^ fragment along with the mouse *Il1b* peak5 and peak6 fragments were PCR amplified and cloned into the XhoI and HindIII sites in the Firefly luciferase plasmid pGL4.23 (Promega) using the Cold Fusion kit (System Biosciences Inc) according to manufacturer instructions. These enhancer-containing plasmids along with pcDNA 3.1-empty vector, pCDNA3.1-Brd4-HA^46^, GFP-RelA (Plasmid #23255, Addgene), and a Renilla luciferase control plasmid were prepared using the HiSpeed Plasmid Maxi kit (Qiagen). Luciferase assays were conducted using the Dual-Luciferase Reporter Assay System (Promega). HeLa cells were seeded at 70,000 cells per well of a 24-well plate approximately 24 hours prior to transfection. All enhancer plasmids (200 ng) were co-transfected with Renilla control plasmid (20 ng) and either a GFP-RelA (400ng), pCDNA3.1-Brd4-HA (400ng), GFP-RelA (200ng) and pCDNA3.1-Brd4-HA (200ng), or empty vector control (400ng) with 2.4 ul Fugene HD (Promega, E2311) in 43 ul Opti-MEM media (Gibco). Each construct was transfected in three technical replicates and at least 3 biological replicates were performed. Luciferase assays were performed 36 hours after transfection according to manufacturer instructions. Measurements were taken using SpectraMax MiniMax 300 imaging cytometer with SoftMax Pro6, Version 6.4 (Molecular Devices). Firefly luciferase activity was normalized with Renilla luciferase activity.

### siRNA transfection of iPSC-FBs

Approximately 8×10^4^ iPSC-CFs were seeded in a 6-well plate the day prior to transfection. Wells that were to be simultaneously stimulated with TGFB/IL1B on the day of transfection were switched to FibroGRO^™^ Basal Medium with 0.5% FBS while the unstimulated wells were switched to FibroGRO^™^ Basal Medium with 2% FBS. Cells were transfected with a final concentration of 15 nM siRNA (Sigma, see details in relevant “Tables” section) using 7ul of Lipofectamine^™^ RNAiMAX transfection reagent (ThermoFisher Scientific) for each well according to the manufacturer instructions. 24 hours after transfection, appropriate wells were treated with TGFB, IL1B, or TGFB+IL1B. Cells were harvested 48-72 hours after stimulation. For RAW 264.7 cells, 8×10^4^ were seeded in 6-well plates the day prior to transfection. Cells were transfected with either 15 nM mouse siRNA (ThermoFisher Scientific) or 15 nM siRNA Control using 7 ul of Lipofectamine^™^ RNAiMAX transfection reagent (ThermoFisher Scientific) for each well according to the manufacturer instructions. Approximately 24 hours after transfection, LPS stimulation was performed and cells were harvested 6 hours after treatment.

### Isolation and culture of mouse bone marrow-derived macrophages (BMDM)

BMDM were isolated from 8-10 week-old male and female heterozygous *Cx3cr1^GFP^* knock-in mice (B6.129P2(Cg)-Cx3cr1tm1Litt/J, Jackson Labs #005582) as previously described^47^. Briefly, animals were anesthetized with isoflurane inhalation to effect. Tibias and femurs were removed, cannulated with a 25g needle, and flushed with sterile DMEM (Fisher Scientific # SH30022FS) containing 10% bovine growth serum (BGS; Fisher Scientific #SH3054103) and 1% penicillin-streptomycin (P/S; VWR #10220-718) to isolate whole bone marrow cells. 4×10^5^ cells were plated and cultured on a 10 cm uncoated Petri plate in DMEM containing 10% bovine growth serum, 1% penicillin-streptomycin, and 50 ng/mL macrophage colony stimulating factor (MCSF; ProSpecBio #CYT-439). A 50% media refresh was performed after 3 days and cells were then split onto assay plates after reaching 70% confluency (days 7-8 post-plating).

### LPS stimulation BMDM for conditioned media experiments and gene expression analysis

Passage #1 BMDM grown and differentiated as above were plated overnight on 6-well plates at 0.5×10^6^ cells per well in DMEM containing 10% BGS, 1% P/S and 50 ng/mL MCSF. Cells were then equilibrated for 4 hrs by changing media to low-serum DMEM containing 1% BGS, 1% P/S and no MCSF. BMDM were then treated for 24 hrs with 100 ng/mL lipopolysaccharide (LPS; Invivogen #tlrl-3pelps), followed by addition of 2mM ATP along with 10 ng/mL of either anti-mouse/rat IL-1β neutralizing antibody (Invivomab #BE0246, Lot #745620M2) or IgG control antibody (Invivomab #BE0091, Lot #765821F1) for an additional 2 hrs. Media was then collected, centrifuged at 500x*g* for 10 min. at 4°C to remove cell debris, and flash-frozen at −80°C. To assess transcriptional upregulation of *Il1b* following LPS stimulation in BMDM, passage #1 BMDMs were treated with 100 ng/mL LPS as above. Cells were lysed in 500 uL of TRIzol (Thermo Scientific #15596018 and RNA was extracted via phase separation with bromo-chloro-propane. cDNA was synthesized using a Verso cDNA Synthesis Kit (Thermo Scientific #AB1453B) and semi-quantitative RT-PCR was performed using SYBR Green (Thermo Scientific #A46012) on an Applied Biosystems StepOne Plus thermocycler.

### Collagen gel contraction assay

Compressible collagen matrices were prepared in 24-well plates using PureCol EZ Gel Solution (Advanced BioMatrix #5074) by incubating at 37°C for 1.5 hrs. Induced pluripotent stem cell-derived cardiac fibroblasts (iPSC-CFs) suspended in serum-supplemented growth serum (FibroGRO^™^ Basal Medium [Millipore #SCMF-BM] with 2% Hyclone^™^ Fetal Bovine Serum [FBS; Fisher #SH30071.02]) were seeded (25,000 cells/gel) on the collagen gels for 24 hrs. Cells were either transferred to low-serum media (FibroGRO^™^ with 0.5% FBS) and stimulated with 5 ng/mL Transforming Growth Factor-β (TGFB; Fisher #50-725-143) and 10 ng/mL Interleukin-1β (IL1B), or alternatively, treated with undiluted BMDM conditioned media described above. Collagen gels were released from the walls of the well, and images were acquired every 24 hrs for up to 120 hrs. Gel area for each well was determined using ImageJ software and data are reported as percent contraction.

### Immunofluorescence Staining

iPSC-CF were seeded in 12-well plates containing glass coverslips at a density of 36,000 cells/well for 24 hrs in FibroGRO^™^ with 2% FBS. Cells were serum-starved in FibroGRO^™^ with 0.5% FBS and simultaneously stimulated with 5 ng/mL TGFB and 10ng/mL IL1B for 72 hrs. Cells were fixed in 3.7% EM-grade PFA in PBS for 10 mins. Fixed cells were permeabilized in 0.25% Triton-X100 and blocked in PBS containing 0.1% Tween-20, 1% bovine serum albumin (BSA; Akron #AK8905) and 22.52 mg/mL glycine for 30 mins. Cells were incubated with a primary antibody directed against smooth muscle α-actin (Abcam #ab7817, 1:100) overnight at 4C. Cells were washed and incubated with Alexa Fluor-conjugated secondary antibody (ThermoFisher) for 1 hr in the dark with 300nM 4’,6-diamidino-2-phenylindole (DAPI). Coverslips were mounted using ProLong^™^ Diamond Antifade Mountant and allowed to cure overnight prior to imaging on a Keyence BZ-X710 fluorescence microscope.

### Bulk RNA sequencing on iPSC-CFs

Total RNA from iPSC-CFs was quantified with a Nanodrop (Thermo scientific). After RNA quality control was determined using an Agilent 2100 bioanalyzer (Agilent Technologies), samples were sent to Novogene Co for sequencing. Samples were sequenced with PE150 on a NovaSeq instrument (Illumina). Bulk RNA-seq raw reads were aligned to the GRCm37 (hg19) genome using the Rsubread aligner^48^. Rsubread featureCounts was used to quantify the number of reads per feature of Ensembl hg19 gene annotation file. For differential gene expression analysis, genes with <2 counts per million (CPM) in less than 50% of samples were removed. For analyses using all genes, no cpm filter was applied. EdgeR^49^ calcNormFactors was used to calculate library size normalization factors. The Limma^50^ “voom” function was used to log2 transform and quantile normalize the CPM matrix. Gene count data was processed to log2 counts per million using the Bioconductor package edgeR (v3.32.1), differential Gene Expression analysis was performed using Limma (v3.46.0)

### *In vivo* antibody-mediated neutralization of IL1B

C57BL/6J mice (Jackson Laboratory, Stock No: 000664) were used to perform TAC and treated with either 500ug of polyclonal Armenian hamster IgG isotype control (Invivomab #BE0091, Lot #765821F1) or anti-IL1B antibody (Invivomab #BE0246, Lot #745620M2) injected intraperitoneally every 3 days for 30 days, starting from the day of the TAC surgeries. At day 30 post-TAC the hearts were isolated and Langendorff perfusion performed. Non-CM cells from both cardiac ventricles were processed for subsequent scRNA-seq.

### Statistics and reproducibility

Standard statistical analyses were performed using GraphPad Prism 9. In all figures, the exact p-value is indicated if the p-value is higher than 0.0001. For values lower than 0.001 we indicate P<0.0001. Figure legends include the information of which specific statistical test has been run to calculate significance. In all panels related to cardiac function, gene expression by RT-qPCR, collagen contraction, and aSMA quantification, the means ± SEM are reported in the figures and the number of replicates is indicated as data points in the graphs.

## Tables

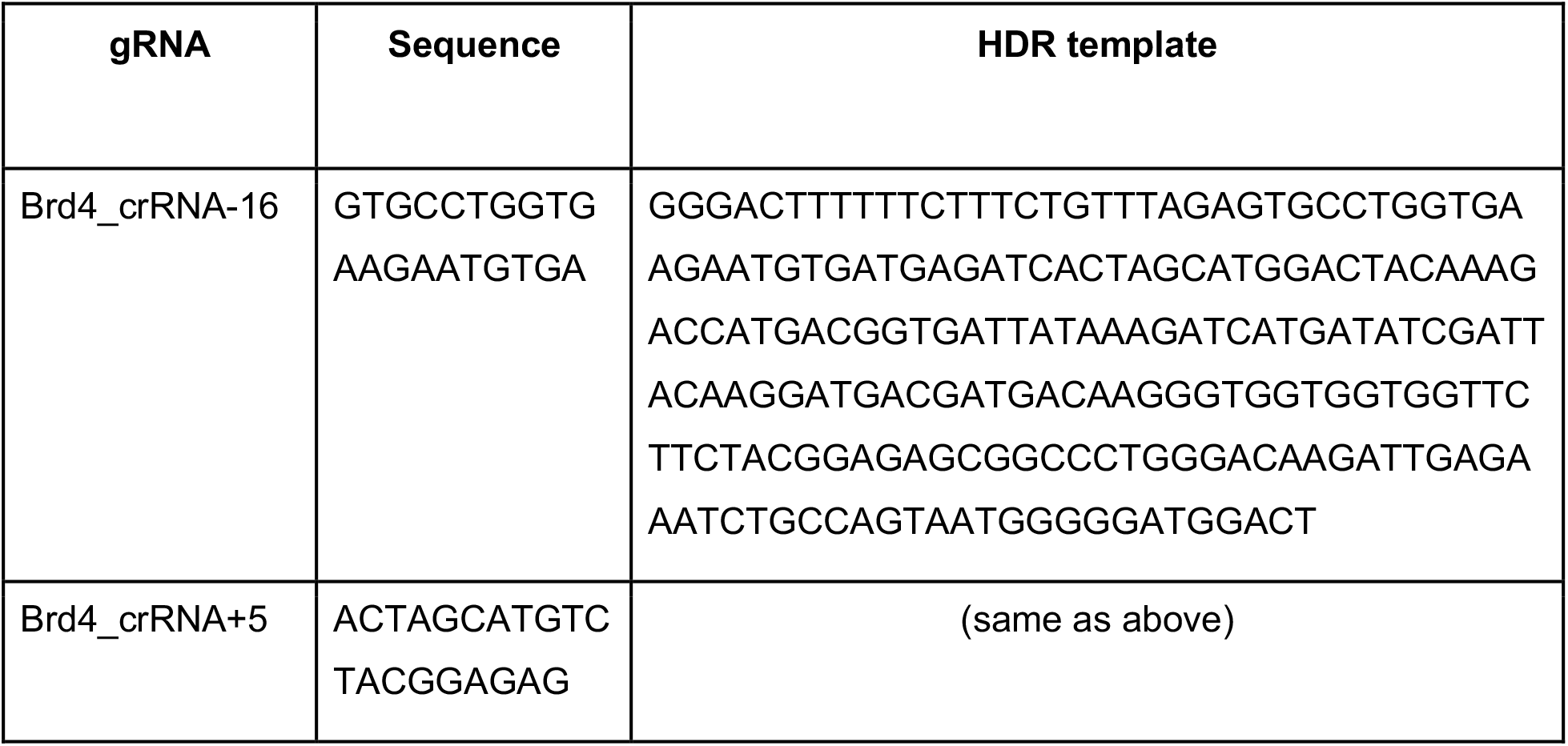
CRISPR rRNA and HDR template to generate Brd4^flag/flag^ mouse.

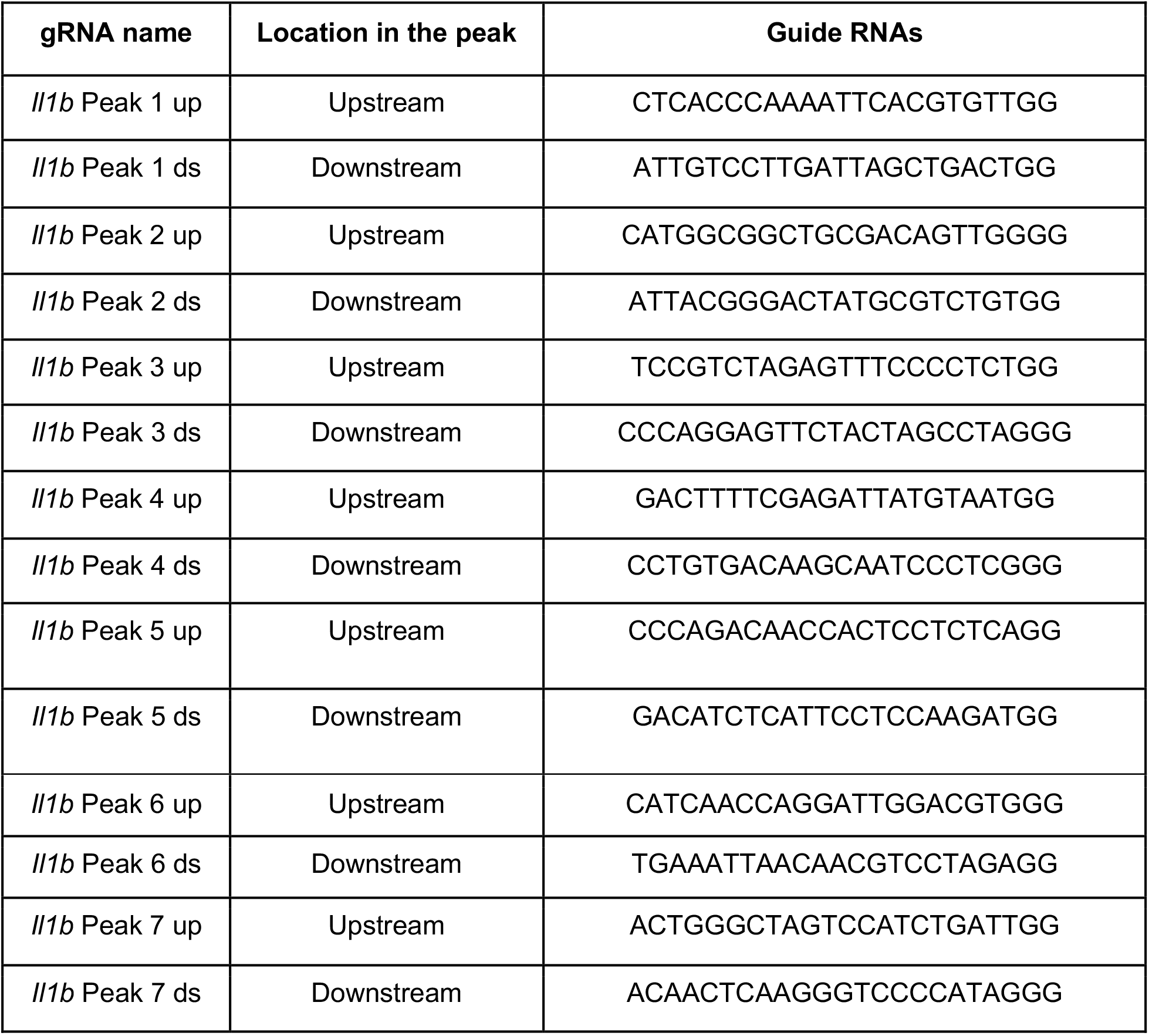
CRISPR guide RNAs to generate *Il1b* enhancer KO clones.

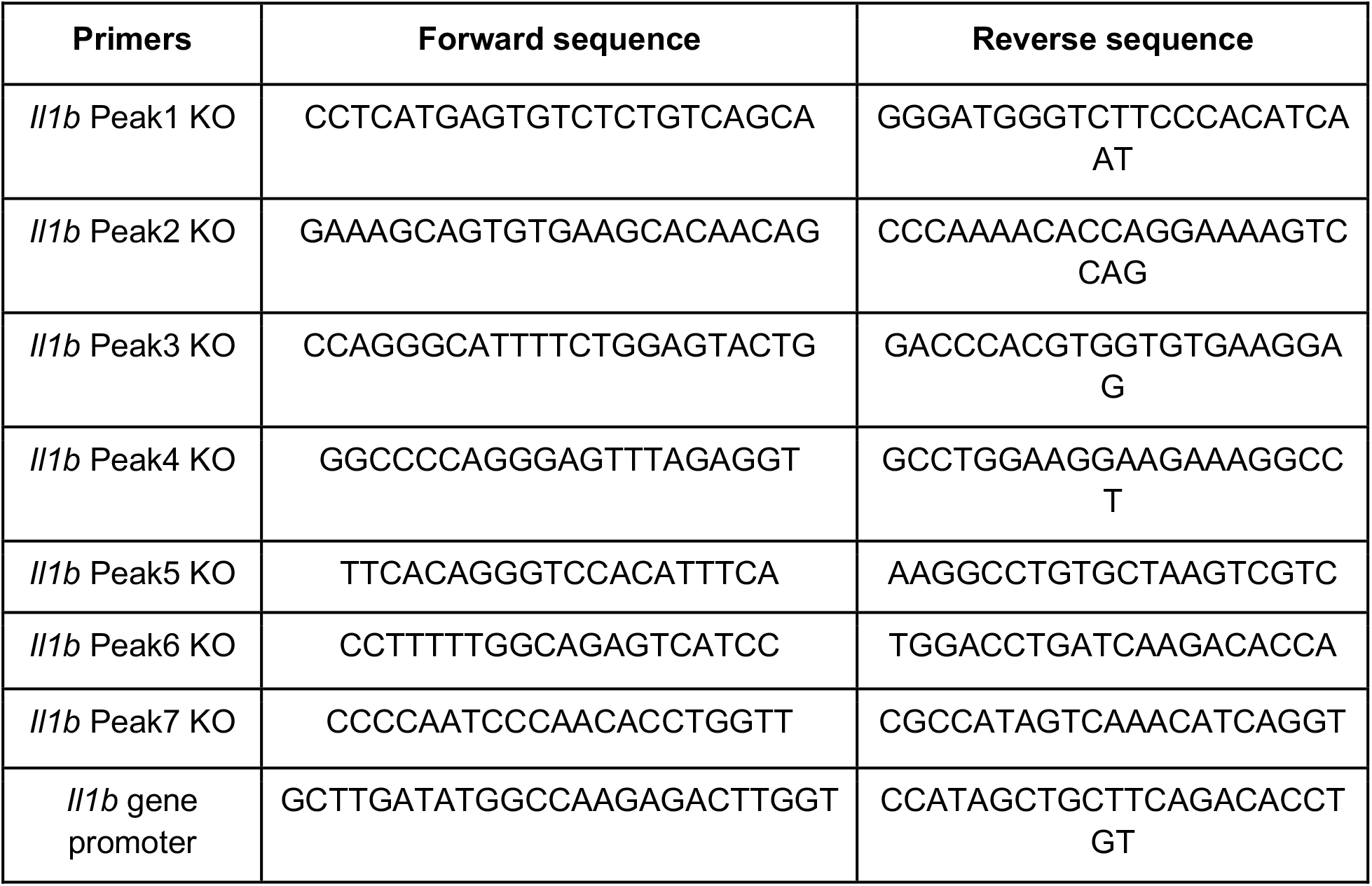
PCR primers to genotype *Il1b* enhancer KO clones.

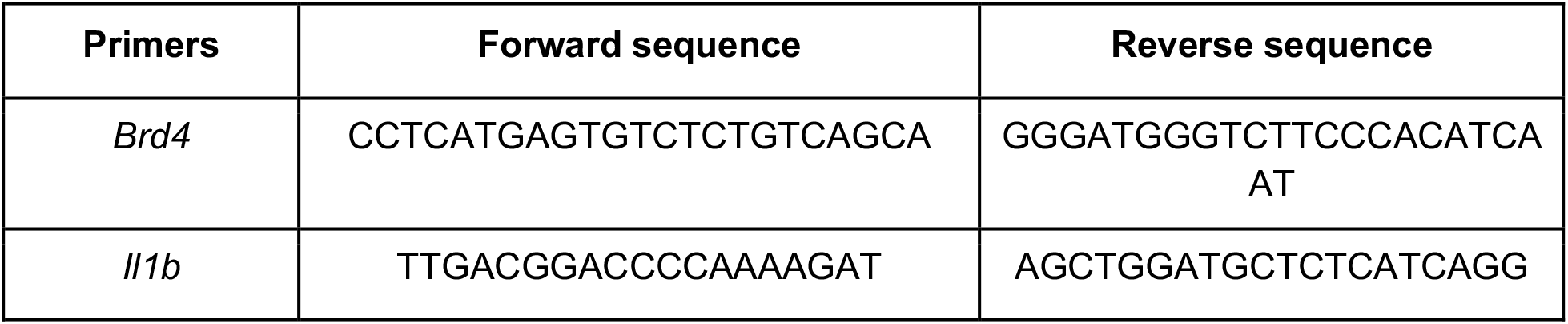
qPCR SYBR probes.

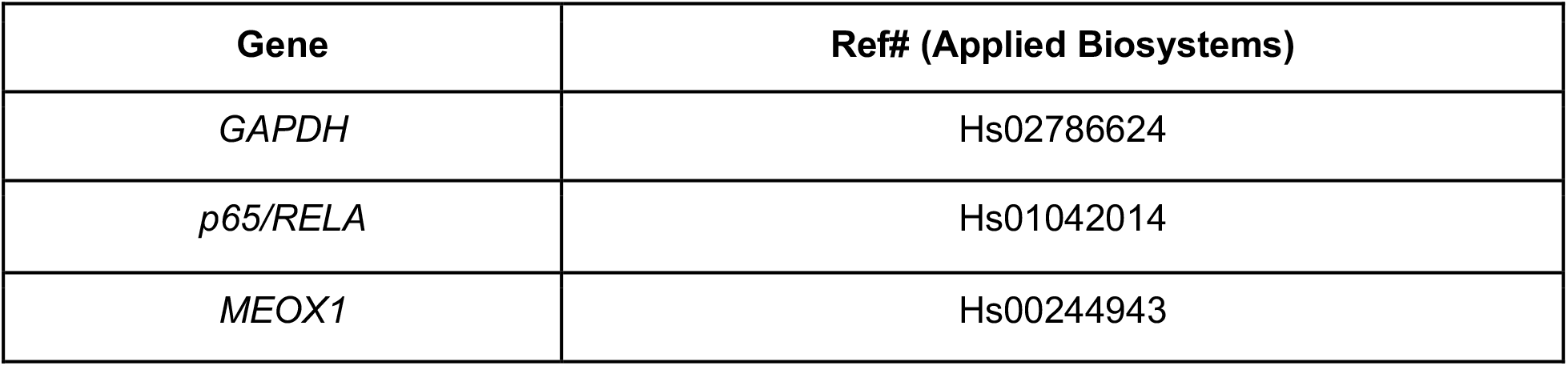
qPCR Taqman probes Taqman probes.

**qPCR Taqman probes Taqman probes**
| **Gene** | **Ref# (Applied Biosystems)** |
| --- | --- |
| *GAPDH* | Hs02786624 |
| *p65/RELA* | Hs01042014 |
| *MEOX1* | Hs00244943 |

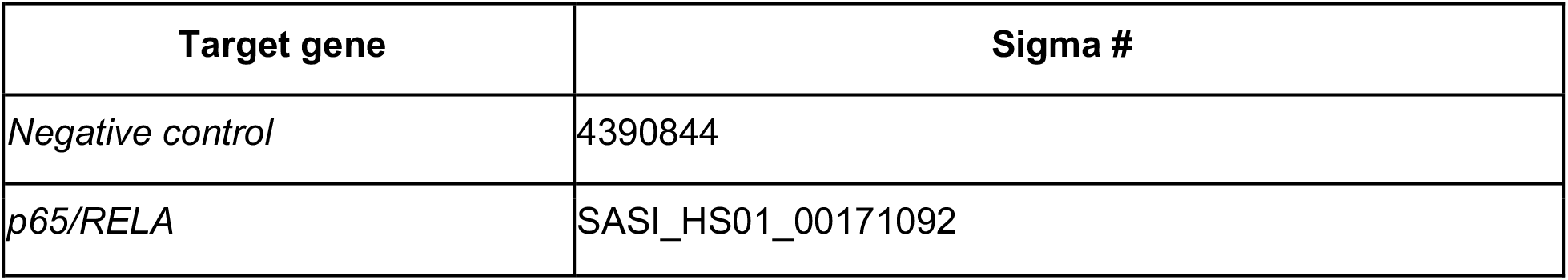
siRNAs.

**siRNAs**
| **Target gene** | **Sigma #** |
| --- | --- |
| *Negative control* | 4390844 |
| *p65/RELA* | SASI_HS01_00171092 |

